# An interaction hub on Ndc80 complex facilitates dynamic recruitment of Mps1 to yeast kinetochores to promote accurate chromosome segregation

**DOI:** 10.1101/2023.11.07.566082

**Authors:** Emily J. Parnell, Erin Jenson, Matthew P. Miller

**Author notes:** Correspondence (M.P.M.).

## Abstract

Accurate chromosome segregation relies on kinetochores carrying out multiple functions, including establishing and maintaining microtubule attachments, forming precise bioriented attachments between sister chromatids, and activating the spindle assembly checkpoint. Central to these processes is the highly conserved Ndc80 complex. This kinetochore subcomplex interacts directly with microtubules, but also serves as a critical platform for recruiting kinetochore-associated factors and as a key substrate for error correction kinases. The precise manner in which these kinetochore factors interact, and regulate each other’s function, remains unknown – considerably hindering our understanding of how Ndc80 complex-dependent processes function together to orchestrate accurate chromosome segregation. Here, we aimed to uncover the role of Nuf2’s CH domain, a component of the Ndc80 complex, in ensuring accurate chromosome segregation. Through extensive mutational analysis, we identified a conserved “interaction hub” comprising two segments in Nuf2’s CH domain, forming the binding site for Mps1 within the yeast Ndc80 complex. Intriguingly, the interaction between Mps1 and the Ndc80 complex seems to be subject to regulation by competitive binding with other factors. Mutants disrupting this interaction hub exhibit defects in spindle assembly checkpoint function and severe chromosome segregation errors. Significantly, specifically restoring Mps1-Ndc80 complex association rescues these defects. Our findings shed light on the intricate regulation of Ndc80 complex-dependent functions and highlight the essential role of Mps1 in kinetochore biorientation and accurate chromosome segregation.

## INTRODUCTION

The precise segregation of duplicated chromosomes into daughter cells is a fundamental process during cell division. This segregation relies on the interactions between microtubules and kinetochores, which are large protein complexes that assemble on the centromeres of each chromosome (reviewed in ^1^). Accurate chromosome segregation relies on several critical functions performed by kinetochores. First, they must establish and maintain attachments to the tips of microtubules, whose dynamic growing and shrinking generates the forces to physically move chromosomes within the cell. Second, kinetochores on each pair of replicated sister chromatids must form bioriented attachments to microtubules originating from opposite spindle poles. Since initial kinetochore-microtubule attachments are randomly established, these attachments are often erroneous and must be detected and corrected to prevent mis-segregation. Third, kinetochores must activate the spindle assembly checkpoint when microtubule attachments are absent, thereby halting the cell cycle until these errors are corrected (reviewed in ^2,3^). The precise mechanisms by which kinetochores coordinate and regulate these diverse functions remain important and open questions in the field.

At the core of these essential cellular functions lies the Ndc80 complex, a highly conserved subcomplex of the kinetochore. This complex is a heterotetrameric, rod-like assembly, comprised of four distinct proteins: Ndc80, Nuf2, Spc24, and Spc25^4–9^. The complex engages with microtubules through multiple domains, including the globular CH-domains located at the N-terminal end of the Ndc80:Nuf2 complex, which provides a positively charged interaction surface for microtubules^7,10,11^. Furthermore, the unstructured, flexible N-terminal tail of the Ndc80 protein harbors an additional microtubule-binding element^7,11–14^. The role of Ndc80’s CH domains in microtubule binding is well-established both in vitro and in vivo^7,10,14,15^. In contrast, the function of Nuf2’s CH domain has remained somewhat enigmatic^7,15^. Structural studies, for instance, indicate that this domain does not make direct contact with the microtubule^6,9,16^. Nevertheless, its high degree of conservation suggests that it likely serves vital functions.

Besides its direct microtubule binding capability, the Ndc80 complex serves as a critical “landing pad” for numerous microtubule-associated proteins, as well as a substrate for various kinases. To enhance the ability to form load-bearing attachments and to track with dynamic microtubule tips, the Ndc80 complex associates with the Dam1 complex in yeast^10,17–22^ and the functionally analogous Ska complex in metazoans^23–26^. The Ndc80 protein itself serves as a critical phosphorylation target for the error correction kinases, namely Ipl1 and Mps1^7,11–13,27–38^. Additionally, the Ndc80 complex directly interacts with additional factors implicated in correcting erroneous attachments, including Stu2/chTOG^39–41^. Lastly, the Ndc80 complex assumes a pivotal role in triggering the spindle assembly checkpoint by recruiting the most upstream component, the kinase Mps1^33–35,37,42–48^. A significant barrier to understanding how these diverse Ndc80 complex-dependent functions are coordinated lies in our limited understanding of how these different factors interact with the Ndc80 complex and the degree to which they influence each other’s functions.

In this study, our aim was to elucidate the role of Nuf2’s CH domain in facilitating chromosome segregation fidelity. Through extensive mutational analysis, we identified a conserved “interaction hub” formed by two segments of Nuf2’s CH domain, its N-terminal loop and G-helix. We demonstrate that this hub constitutes a key portion of the Mps1 binding interface within the yeast Ndc80 complex. This same region of Nuf2 associates with additional factors, including the Dam1 complex, suggesting that Mps1’s kinetochore localization may be regulated by competitive binding. Mutants that disrupt this interaction hub exhibit defects in spindle assembly checkpoint function and severe chromosome segregation errors. Remarkably, Mps1 appears to be the critical factor that binds to this region, since the cellular defects can be rescued by specifically restoring Mps1-Ndc80 complex association. Our work sheds light on the manner in which numerous Ndc80 complex-dependent functions are regulated and demonstrates the essential role of Mps1 in kinetochore biorientation and chromosome segregation.

## RESULTS

### Identification of a Nuf2 patch that is essential for cell viability

For this study, we wished to determine the role that Nuf2’s CH domain plays in the establishment of correctly bioriented sister chromatids. To identify residues that are important for its function, we first looked for spatial patterns of amino acid conservation within Nuf2’s CH domain. Here, we aligned Nuf2 sequences from various fungal species and then mapped the resulting conservation scores onto a previously determined Ndc80 complex structure^49^ (PDB: 5TCS). This analysis revealed a region of conserved residues in close spatial proximity, which is centered on Nuf2’s so called “G-helix” as well as its amino-terminal “loop” (Fig 1A & Fig S1A). These residues, especially F8, P9, R118, S124, N128 and R131, appear highly conserved in fungal and metazoan Nuf2 sequences (Fig 1A & Fig S1A-B). To examine the functional importance of these residues, we generated strains containing an auxin-inducible degron of the endogenous *NUF2*, and ectopically expressed mutant *nuf2* alleles to examine their phenotype in the presence of auxin. We focused on mutating residues that showed strong conservation, and assayed effects on cell viability using a spot dilution assay. We found that many of the mutations resulted in severe cell viability defects, and the most penetrant phenotypes were associated with mutations within the Nuf2 G-helix (*nuf2^S124D^, nuf2^A125D^, nuf2^N128A^* and *nuf2^R131E^*) as well as N-terminal loop (*nuf2^F8A^ ^P9A^*; Fig 1B & Fig S1D). Importantly, these mutations do not appear to affect Nuf2 expression levels (Fig S1C). A summary of all mutants and associated viability phenotypes can be found in Fig 1C and Fig S1D. Based on the severity of viability defects, amount of conservation, and degree of spatial proximity, we chose to focus on mutations *nuf2^F8A^ ^P9A^*, *nuf2^S124D^*, and *nuf2^N128A^* for further analysis.

**Figure 1.**
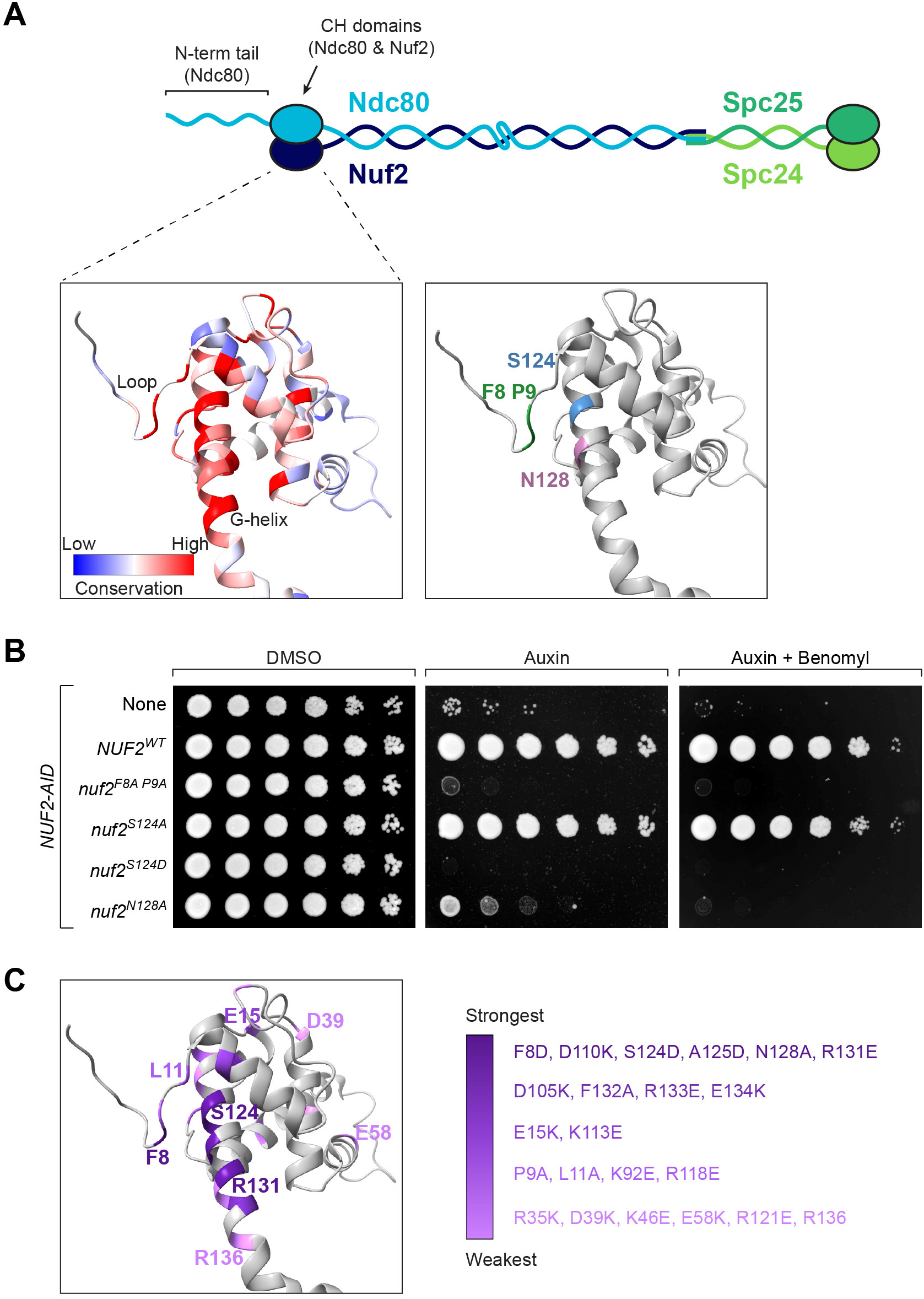
Mutation of conserved Nuf2 residues affects cell viability. (A) Cartoon of the heterotetrameric Ndc80 complex (Ndc80, Nuf2, Spc24, Spc25). CH domains are represented as ovals; coiled-coil domains and the N-terminal tail of Ndc80 are shown as wavy lines. Structure shows *S. cerevisiae* Nuf2 from ^49^ (5TCS), with other members of the Ndc80 complex not shown. Left: Conservation of Nuf2 among 12 fungal species (listed in Fig S1), viewed using ChimeraX, on a scale of -2.5 to +2.5, with the most conserved residues in red and least conserved in blue. Right: Conserved residues used throughout this work are highlighted (F8 and P9 in the N-terminal loop; S124 and N128 in the G-helix). (B) Yeast cell viability assay with a subset of *nuf2* mutant alleles. Strains carry *NUF2-AID* and an ectopic copy of *NUF2-3HA* (No covering allele, “None”, M1889; *NUF2^WT^*, M2038; *nuf2^F8A^ ^P9A^*, M2042; *nuf2^S124A^*, M2040; *nuf2^S124D^*, M2041; *nuf2^N128A^*, M2414). Cells were serially diluted five-fold and spotted onto plates containing DMSO (control), 250 µM auxin (to degrade endogenous Nuf2-AID protein), or 250 µM auxin + 6.5 µg/mL benomyl. (C) Structure of Nuf2 (5TCS), illustrating the range of cell viability phenotypes of mutant alleles observed on plates containing benomyl (data shown in Fig S1D). Nuf2 mutants are listed in groups in decreasing shades of purple, from the strongest phenotype (death on auxin only) to the weakest (slight sickness on auxin + benomyl).

### Spindle assembly checkpoint function is perturbed in nuf2 patch mutants

To begin understanding how these *nuf2* CH domain mutants result in cell viability defects, we first compared the growth phenotypes to previously described mutations in the CH domain of Nuf2’s binding partner, Ndc80. We found that *nuf2* mutants, including *nuf2^F8A^ ^P9A^* and *nuf2^N128A^*, display a considerably more severe viability defect than *ndc80* mutants which perturb microtubule binding^7,10^ (i.e. *ndc80^K204E^* and *ndc80^K122E^ ^K204E^*; Fig 2A and Fig S2). Furthermore, while *ndc80^K122E^ ^K204E^* mutants arrest in mitosis^10^, likely by activating the spindle assembly checkpoint, the *nuf2* mutants did not induce a specific cell cycle arrest (Fig 2B). These observations suggest that these *nuf2* CH domain mutations affect a kinetochore function beyond just microtubule binding. Given the lack of cell cycle arrest, we next assessed spindle assembly checkpoint (SAC) function by examining the localization of the SAC component, Bub1, which is recruited to kinetochores upon SAC activation^50–53^. For this assay, *CDC20-AID* cells were arrested in metaphase by the addition of auxin. Once arrested in metaphase, cells were treated with nocodazole, and co-localization of Bub1-GFP with the kinetochore marker Mtw1-mCherry was monitored. Approximately 72 ± 6% of *NUF2^WT^* cells showed Bub1-GFP recruitment to the kinetochore upon nocodazole treatment, consistent with prior observations^51,53^. In contrast, *nuf2^F8A^ ^P9A^* and *nuf2^N128A^*showed a dramatic decrease in the number of cells with Bub1 at kinetochores (14 ± 5.5% and 14 ± 3.5%, respectively; Fig 2C), consistent with defects in spindle assembly checkpoint function. We also note that Mtw1-mCherry distribution appeared normal in the *nuf2* mutants, suggesting that these mutations do not cause gross defects in kinetochore organization (Fig 2C).

**Figure 2.**
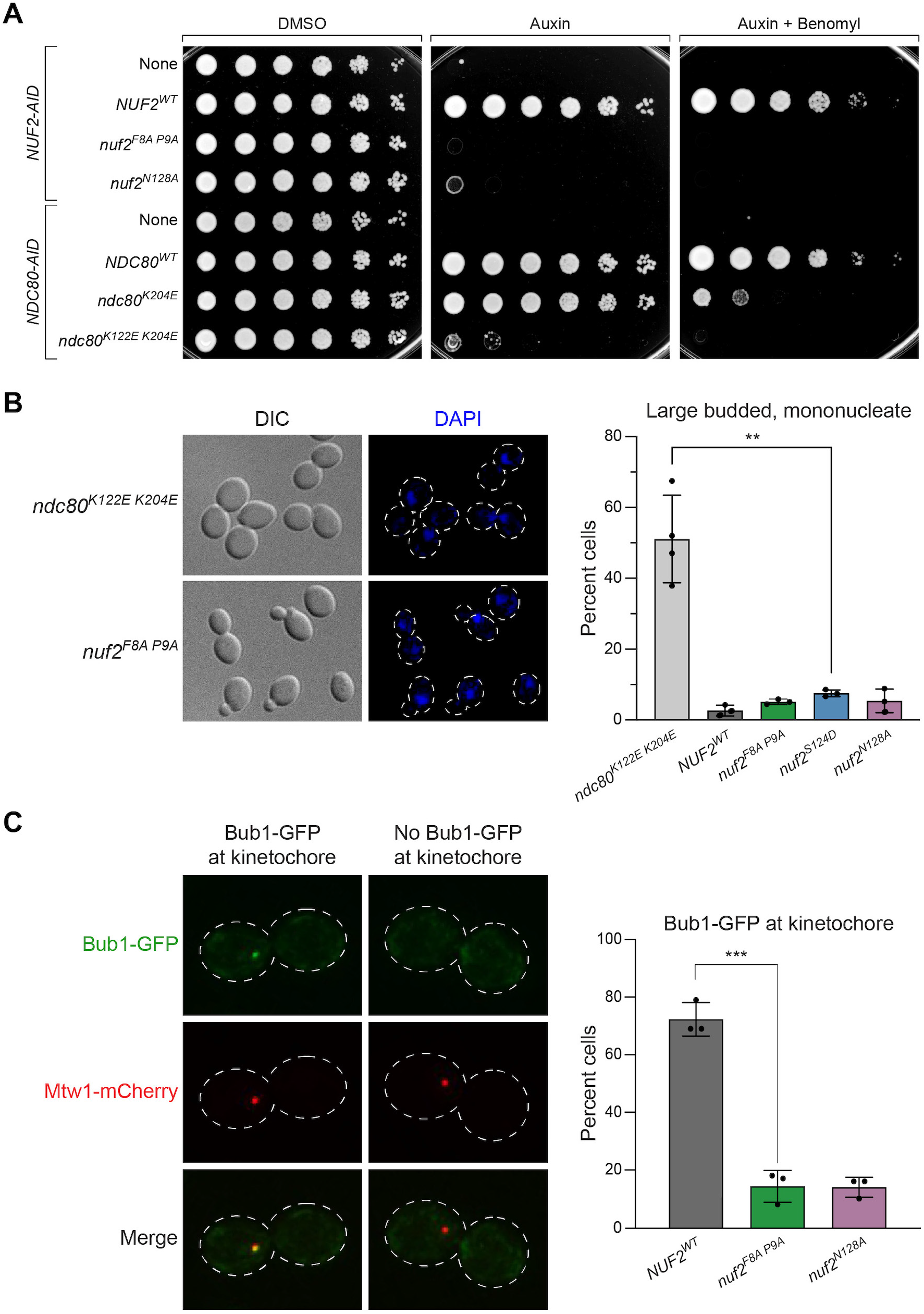
Lethal *nuf2* mutant alleles fail to trigger the spindle assembly checkpoint. (A) Yeast cell viability assay comparing *nuf2* mutants with previously characterized *ndc80* alleles. Strains carry either *NUF2-AID* or *NDC80-AID* at their endogenous loci, and ectopic copies of *NUF2-3HA* (None, M1889; *NUF2^WT^*, M2038; *nuf2^F8A^ ^P9A^*, M2042; *nuf2^N128A^*, M2414) or *NDC80-3HA* (None, M4182; *NDC80^WT^*, M692; *ndc80^K204E^*, M693; *ndc80^K122E^ ^K204E^*, M694). Cells were serially diluted five-fold and spotted onto plates containing DMSO, 250 µM auxin or 250 µM auxin + 6.5 µg/mL benomyl. (B) Comparison of yeast cells expressing *nuf2* mutant alleles to those expressing an *ndc80* mutant allele that induces a metaphase arrest. Exponentially growing strains with *NDC80-AID* or *NUF2-AID* and ectopic copies of either *ndc80^K122E^ ^K204E^-3HA* (M694) or *NUF2-3HA* (*NUF2^WT^*, M2933; *nuf2^F8A^ ^P9A^*, M2935; *nuf2^S124D^*, M2936; *nuf2^N128A^*, M2945) were treated with 500 µM auxin for 2.5 hours to degrade endogenous Ndc80-AID or Nuf2-AID protein prior to fixation. Left: Representative micrographs. Right: Quantitation of cells showing the percentage of large-budded, mononucleate cells in *ndc80* or *nuf2* mutants, with error bars indicating the standard deviation among three replicates (n ≥ 100 cells for each replicate). Significance was determined by a two-tailed unpaired *t* test (**; *P* ≤ 0.01). (C) Bub1-GFP assay for SAC activity. Exponentially growing *NUF2-AID CDC20-AID* strains (*NUF2^WT^*, M2792; *nuf2^F8A^ ^P9A^*, M2794; *nuf2^S124D^*, M2795; *nuf2^N128A^*, M2801) were treated with 500 µM auxin for 2 hours to arrest cells in metaphase, followed by 10 µM nocodazole for 1 hour to activate the SAC. Strains carry *BUB1-GFP* (SAC) and *MTW1-mCherry* (kinetochore) alleles. Left: Representative micrographs. Right: Quantitation of percent of large-budded, mononucleate cells with Bub1-GFP localized at the kinetochore, with error bars indicating the standard deviation among three replicates (n= 66-112 cells for each replicate). Significance was determined by a two-tailed unpaired *t* test (***; *P* ≤ 0.001). Only cells with a single Mtw1 signal, indicating collapse of the spindle, were analyzed.

### Mutants within Nuf2’s patch result in substantial chromosome segregation defects

While these *nuf2* CH domain mutants display spindle assembly checkpoint defects, the lethal phenotypes observed cannot be explained by this limitation alone since the SAC is not essential for cell viability in yeast (Fig 3A). This discrepancy led us to examine sister kinetochore biorientation and chromosome segregation defects as a potential cause of the lethality. To monitor biorientation, we induced a metaphase arrest by depleting Cdc20 (using a methionine-repressible *CDC20* allele) in cells that also carry a fluorescently marked centromere of chromosome III^54^. Under these conditions, opposing spindle pulling forces cause bioriented sister chromatids to separate, appearing as two distinct GFP puncta^49^. The frequency of biorientation in *NUF2^WT^* cells was similar to those typically observed in this assay (44 ± 6%, Fig 3B). However, in *nuf2* mutant cells, we observed a marked reduction in biorientation of CEN III (*nuf2^F8A^ ^P9A^*, 10 ± 1.5%; *nuf2^S124D^*, 6.6 ± 1.5%; and *nuf2^N128A^*, 18 ± 1.5%). Furthermore, we noted a significant portion of *nuf2* mutant cells bypassing the arrest induced by Cdc20 depletion (Fig S3A). In these *nuf2* mutants, spindle elongation led exclusively to instances of sister chromatid non-disjunction and unequal DNA masses (Fig S3B-C). These findings can be attributed to substantial deficiencies in biorientation, where many or all pairs of sister chromatids form syntelic attachments to the mitotic spindle, consequently lacking the necessary forces to restrict spindle elongation.

**Figure 3.**
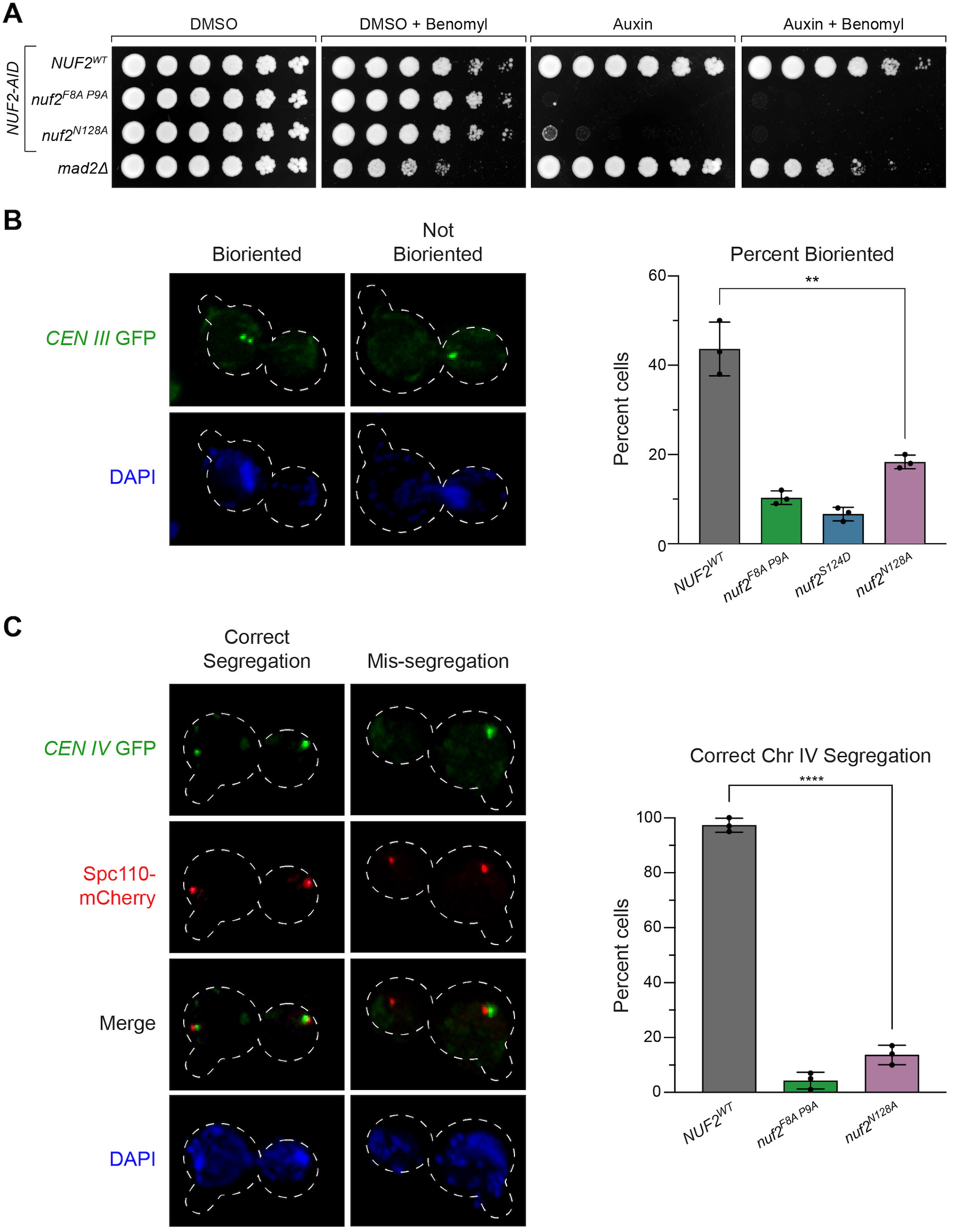
Inviable *nuf2* mutants display a loss of sister chromatid biorientation and mis-segregation of chromosomes. (A) Comparison of *nuf2* mutant cell viability relative to the *mad2* SAC mutant. Strains carrying a *NUF2-AID* and ectopic copies of *NUF2-3HA* (*NUF2^WT^*, M2038; *nuf2^F8A^ ^P9A^*, M2042; *nuf2^N128A^*, M2414) are shown in comparison to *mad2Δ* (M35). Cells were serially diluted five-fold and spotted onto plates containing DMSO, 250 µM auxin, 6.5 µg/mL benomyl or both auxin and benomyl. (B) Assay for sister chromatid biorientation in *nuf2* mutants. Exponentially growing cells carrying *NUF2-AID* and ectopic copies of *NUF2-3HA* (*NUF2^WT^*, M4245; *nuf2^F8A^ ^P9A^*, M4251; *nuf2^S124D^*, M4249; *nuf2^N128A^*, M4253), *pMET-CDC20*, and *CEN III* marked with GFP (*CEN III:lacO LacI-GFP*) were arrested with 1 µg/mL alpha factor for 3 hours, followed by release into fresh media containing methionine (to induce metaphase arrest via *pMET-CDC20*) and 500 µM auxin (to degrade endogenous Nuf2-AID) for 1.5 hours. Left: Representative micrographs. Right: Quantitation of percent of large-budded, mononucleate cells displaying biorientation of the chromosome-marked GFP signal, with error bars indicating the standard deviation among three replicates (n= 71-137 cells for each replicate). Significance was determined by a two-tailed unpaired *t* test (** *P* ≤ 0.01). (C) Chromosome segregation assay in *nuf2* mutants. Exponentially growing cells carrying *NUF2-AID* and ectopic copies of *NUF2-3HA* (*NUF2^WT^*, M4469; *nuf2^F8A^ ^P9A^*, M4475; *nuf2^N128A^*, M4479), as well as *CEN IV* marked with GFP (*CEN IV:lacO LacI-GFP*) and *SPC110-mCherry* (spindle pole marker) were arrested with 1 µg/mL alpha factor for 3 hours, followed by release into fresh media containing 500 µM auxin for 2 hours. Left: Representative micrographs. Right: Quantitation of percent of binucleate cells displaying correct segregation of the chromosome-marked GFP signal, with error bars indicating the standard deviation among three replicates (n = 75-151 cells for each replicate). Significance was determined by a two-tailed unpaired *t* test (****; *P* ≤ 0.0001).

To examine chromosome segregation fidelity in these *nuf2* mutants, cells again carrying a fluorescently marked centromere (now of chromosome IV) were released from a G1 arrest, and chromosome segregation was examined upon anaphase onset. Cells expressing *nuf2^F8A^ ^P9A^*and *nuf2^N128A^* displayed surprisingly high errors in segregating chromosome IV (*NUF2^WT^* 96 ± 4.6% correct segregation compared to *nuf2^F8A^ ^P9A^* 4 ± 3.1%, *nuf2^N128A^* 14 ± 3.5%, Fig 3C and Fig S3D). Together, these observations show that the region of Nuf2’s CH domain, consisting of its G-helix and N-terminal loop, are required for spindle assembly checkpoint function and essential for biorientation and accurate chromosome segregation.

### Mps1-kinetochore localization perturbed in nuf2 patch mutants

The observed defects in both SAC function and biorientation led us to consider the idea that decreased recruitment of the Mps1 kinase to the kinetochore may be the cause of the observed phenotypes. Mps1 lies at the apex of spindle assembly checkpoint signaling. Beyond this, its activity is important in the establishment of correctly bioriented kinetochore attachments^32–37,44,45^, and Nuf2 and Ndc80 have been implicated as the binding site for Mps1^30,42,43,55^. We therefore used an established kinetochore co-IP assay to examine the levels of Mps1 associated with the kinetochore in *NUF2^WT^* relative to *nuf2* mutants^31,56^. Kinetochores were immunoprecipitated via anti-Flag pulldown of the kinetochore component Dsn1-Flag, and co-purification of Mps1-3V5 was determined by western blot. As expected, *NUF2-AID* cells (treated with auxin 2 hours prior to harvesting) expressing *NUF2^WT^* showed Mps1 co-purification with kinetochores. In contrast, *nuf2^F8A^ ^P9A^*, *nuf2^S124D^*, and *nuf2^N128A^* mutants displayed nearly undetectable Mps1 binding but preserved Ndc80 association (Fig 4A). We propose this region of Nuf2 forms a key part of the Mps1 binding interface to the yeast Ndc80 complex. These results are also consistent with Yu and colleagues who made a similar observation for human Mps1 binding to Ndc80 complex^43^, suggesting this is a remarkably conserved interface.

**Figure 4.**
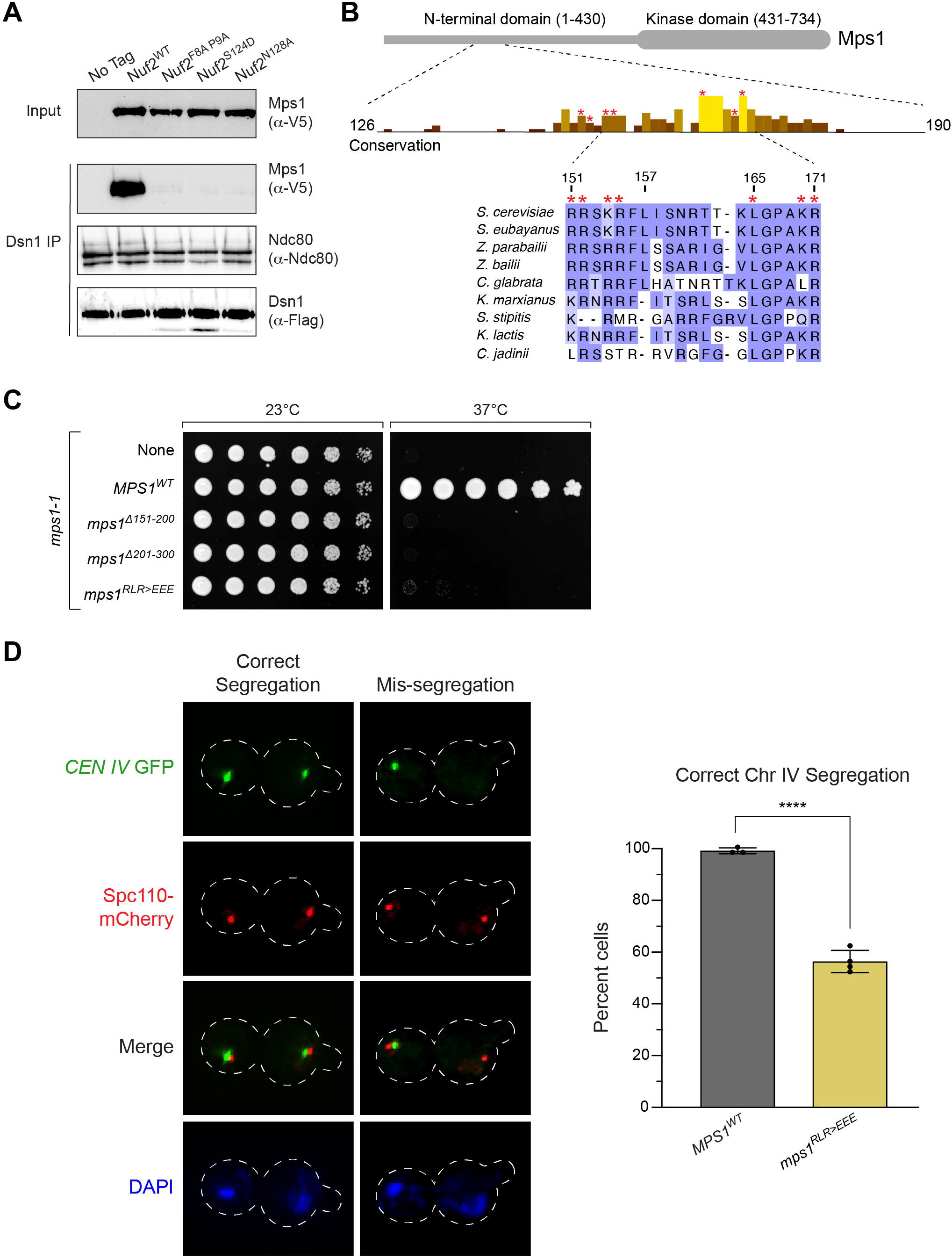
Mps1 kinetochore recruitment is lost in *nuf2* mutants, and an *mps1* mutant partially phenocopies a *nuf2* mutant for chromosome segregation defects. (A) Detection of Mps1-3V5 association with the kinetochore via immunoprecipitation. Exponentially growing cultures expressing *DSN1-6His-3Flag* (No tag, M2630) as well as *MPS1-3V5, NUF2-AID* and ectopic copies of *NUF2-3HA* (*NUF2^WT^*, M3494; *nuf2^F8A^ ^P9A^*, M3500; *nuf2^S124D^*, M3498; *nuf2^N128A^*, M3504), were treated with auxin 2 hours prior to harvest. Kinetochores were purified from lysates by anti-Flag immunoprecipitation (IP) and analyzed by immunoblotting. (B) Cartoon of the Mps1 protein and a conservation histogram for amino acids 126 to 190. The region from 151 to 171 is shown in detail, with amino acids mutated in this work marked by an *. Numbering is according to the *S. cerevisiae* sequence. (C) Complementation of the temperature sensitive *mps1-1* allele (M56) by ectopically expressed *MPS1* (*MPS1^WT^*, M4407; *mps1^Δ151–200^*, M4412; *mps1^Δ201–300^*, M4413; *mps1^RLR>EEE^*, M4949). Cells were serially diluted five-fold, spotted onto plates, and grown at *mps1-1* permissive (23°C) and non-permissive (37°C) temperatures. (D) Chromosome segregation assay in *mps1* mutants. Exponentially growing cells carrying the *mps1-1* temperature sensitive allele and ectopic copies of *MPS1* (*MPS1^WT^*, M4714; *mps1^RLR>EEE^*, M4710, M4947, M4948), as well as *CEN IV* marked with GFP (*CEN IV:lacO LacI-GFP*) and *SPC110-mCherry* (spindle pole marker), were arrested with 1 µg/mL alpha factor for 3 hours, followed by release into fresh media at 37°C for 2 hours. Due to the difference in phenotypes between *nuf2* mutants and *mps1^RLR>EEE^*, three independent strains were analyzed to ensure validity of the result. Left: Representative micrographs. Right: Quantitation of percent of binucleate cells displaying correct segregation of the chromosome-marked GFP signal, with error bars indicating the standard deviation among three to four replicates (n = 61-154 cells for each replicate). Significance was determined by a two-tailed unpaired *t* test (****; *P* ≤ 0.0001).

If the *nuf2* mutant defects stem from reduced Mps1 binding, we hypothesized that an *mps1* mutant at this interface would exhibit a similar phenotype. To identify potential interaction regions of Mps1 with the Ndc80 complex, we sought conserved regions within Mps1 outside its C-terminal kinase domain and identified two highly conserved patches encompassing residues 151-157 and 165-171 (Fig 4B). Earlier studies associated residues 151-200 with Mps1’s SAC and biorientation functions, while finding residues 201-300 are crucial for spindle pole body duplication^57^. To assess the functional significance of these conserved Mps1 segments, we initially examined cell viability. Here, we used a temperature sensitive allele, *mps1-1*, to disrupt the endogenous function, as an *MPS1-AID* allele did not fully disrupt cell viability in our strain background (data not shown). Consistent with previous observations, *mps1^Δ151–200^* and *mps1^Δ201–300^* mutants were inviable in this system^57^ (Fig 4C). While neither *mps1^Δ151–157^*nor *mps1^Δ165–171^* fully disrupted cell viability, combining these deletions or introducing point mutations spanning one or both regions resulted in complete inviability (Fig 4C and Fig S4A). The notion that this portion of Mps1 facilitates binding to the Ndc80 complex is well-supported by concurrent studies from Pleuger *et al.*^58^ and Zahm *et al.*, employing biochemical and structural approaches to determine the Mps1-Ndc80 complex interface (personal communication S. Westermann and S. Harrison).

Based on the above observations we decided to further characterize the *mps1^R155E^ ^L165E^ ^R170E^* mutant (hereafter designated as “*mps1^RLR>EEE^*”), focusing on spindle pole body separation and chromosome segregation. Cells carrying a fluorescently marked centromere of chromosome IV were arrested in G1 at a permissive temperature. Upon release from this G1 arrest at a non-permissive temperature, we examined spindle pole body separation and chromosome segregation during anaphase. *mps1^RLR>EEE^* cells showed spindle pole body separation similar to that of *MPS1^WT^*, indicating this role of Mps1^59^ remains functional in these mutants (Fig S4B). In contrast, *mps1^RLR>EEE^*mutants displayed a large degree of chromosome segregation errors (*MPS1^WT^*99 ± 1.4% correct segregation, compared to *mps1^RLR>EEE^* 56 ± 4.3%, Fig 4D and Fig S4C). It is worth noting that the chromosome segregation defects observed in the *mps1^RLR>EEE^* mutant were not as pronounced as those observed in the *nuf2* CH domain mutants (14% correct segregation for *nuf2^N128A^*, compared to 56% for *mps1^RLR>EEE^*, Fig 3C and Fig 4D). This discrepancy may be due to the *mps1^RLR>EEE^* mutant not completely disrupting Mps1’s ability to bind the Ndc80 complex. Alternatively, it is plausible that additional factors might bind this region of Nuf2’s CH domain, and the observed phenotypes could arise from the combination of these defects.

### Dam1 complex recruitment to kinetochores is altered in nuf2 CH domain mutants

The inconsistency in observed chromosome segregation defects between the *nuf2* and *mps1* mutants prompted us to consider that the *nuf2* mutants might disrupt binding to additional factors. Recent structural studies have suggested this region of the Ndc80 complex as a potential interaction site for the Dam1 complex^19,20^. To examine if the *nuf2* CH domain mutants disrupt Dam1 recruitment to the kinetochore, we assessed the localization of a component of the Dam1 complex, Ask1-YFP, along the mitotic spindle in metaphase arrested cells. *NUF2^WT^* cells exhibited a characteristic bilobed distribution of Ask1 signals proximal to the spindle pole (marked by Spc110-mCherry), consistent with expected kinetochore localization. On the other hand, *nuf2^F8A^ ^P9A^* mutant cells displayed a substantial change in Ask1 localization, showing a dramatic decrease proximal to the spindle pole, and a noticeable increase in localization along the mitotic spindle (Fig 5A). These observations align with the *nuf2^F8A^ ^P9A^* mutant disrupting normal localization of the Dam1 complex to the kinetochore. If Dam1 complex localization is compromised in *nuf2* mutants, we might anticipate synthetic lethality with hypomorphic alleles of Dam1 complex components. In line with this idea, we found that *nuf2^F8A^*, *nuf2^P9A^*, and *nuf2^I10A^ ^L11A^* display synthetic lethality and/or sickness with *dad1-1* mutants (Fig 5B and Fig S5). Collectively, these findings support the notion that multiple factors, including Mps1 and the Dam1 complex, interact with this “interaction hub” of Nuf2’s CH domain, and the observed phenotypes are likely a result of the combined effects of these binding alterations. This also raises an intriguing possibility that competition for binding to the Ndc80 complex might contribute to regulating the localization and function of these factors.

**Figure 5.**
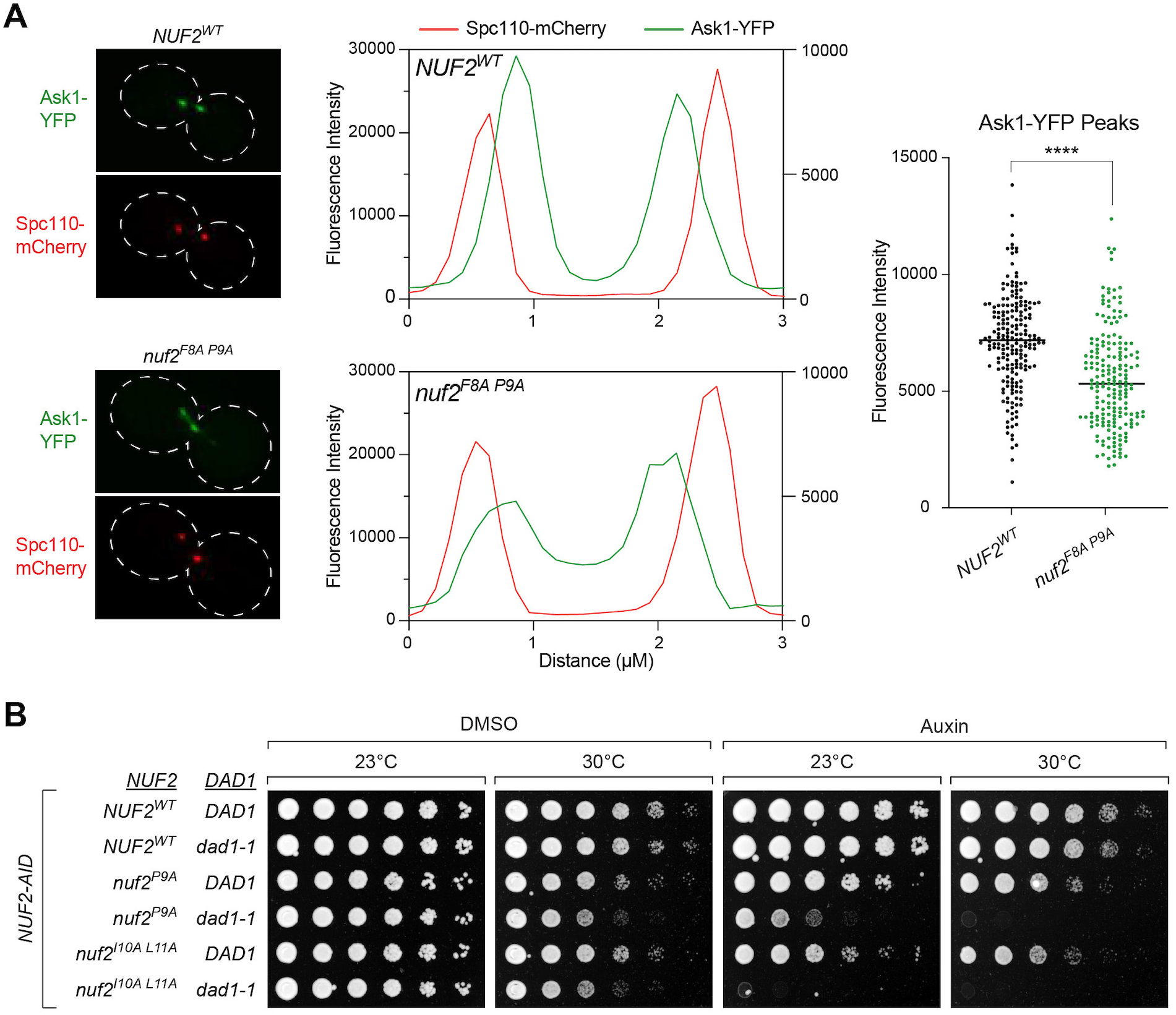
Dam1 localization to the kinetochore is disrupted in *nuf2* mutants. (A) Ask1-YFP localization assay in *NUF2^WT^* and *nuf2^F8A^ ^P9A^*. Strains carry a *NUF2-AID* allele and ectopic copies of *NUF2-3HA* (*NUF2^WT^*, M4985; *nuf2^F8A^ ^P9A^*, M4987), as well as *CDC20-AID*, *ASK1-YFP* (Dam1 complex) and *SPC110-mCherry* (spindle pole marker). Left: Representative micrographs. Middle: Profile plots showing intensities of a line scan through the spindles of *NUF2^WT^* and *nuf2^F8A^ ^P9A^* cells. The Spc110-mCherry signal intensity (red) was plotted on the left y-axis, and the Ask1-YFP signal intensity (green) was plotted on the right y-axis. Right: Graph of YFP intensities, with each dot representing an individual spindle pole-proximal peak (n ≥ 100 cells, ≥ 200 peaks each for *NUF2^WT^* and *nuf2^F8A^ ^P9A^*, respectively). Significance was determined by a two-tailed unpaired *t* test (****; *P* < 0.0001). (B) Cell viability assay assessing synthetic lethality of *nuf2* mutant alleles with the temperature sensitive *dad1-1* allele. Strains carry a *NUF2-AID* allele and ectopic copies of *NUF2-3HA* (*NUF2^WT^ DAD1*, M2038; *NUF2^WT^ dad1-1*, M4465; *nuf2^P9A^ DAD1*, M2152; *nuf2^P9A^ dad1-1*, M4926; *nuf2^I10A^ ^L11A^ DAD1*, M3750; *nuf2^I10A^ ^L11A^ dad1-1*, M4928). Cells were serially diluted five-fold, spotted onto plates containing DMSO or 250 µM auxin and grown at *dad1-1* permissive (23°C) and semi-permissive (30°C) temperatures.

### Ndc80 complex interaction hub mutants can be suppressed by modulating Mps1 levels

Considering that multiple factors seem to interact with this interaction hub, such as Mps1, the Dam1 complex, and potentially other factors, including the Aurora B kinase, Ipl1 (see Zahm *et al.*, personal communication with S. Harrison), a challenging question arises: how do we determine the relative importance of each of these factors in promoting accurate chromosome segregation? Some insight into this question came from the following observations. We reasoned that if defects in Mps1-kinetochore association was the main cause of viability defects in our *nuf2* mutants, perhaps modulating Mps1 levels at the kinetochore could suppress the associated defects. Prior research with human proteins demonstrated that the presence of the N-terminal tail of Ndc80 diminishes Mps1’s capacity to bind to the Ndc80 complex in vitro, likely due to competitive binding. Furthermore, this inhibition appears to be relieved by phosphorylation of the N-terminal tail^43^. If this idea is correct, and Ndc80’s N-terminal tail forms a ‘closed’ conformation to compete with Mps1’s ability to bind in cells, we reasoned that deletion of this tail should increase kinetochore-associated Mps1 (Fig 6A) and may suppress the *nuf2* mutant growth defects. We constructed a *NUF2-AID NDC80-AID* ‘double AID’ strain to conduct these experiments, and found that this genetic background was far more sensitized, *i.e. nuf2* mutants which show no detectable growth defect in a *NUF2-AID* ‘single AID’ background, now result in dramatic cell viability defects (Fig S6A). Consistent with the idea above, many of the *nuf2* mutants at the described Mps1 binding interface showed prominent growth defects in the context of *NDC80^WT^*, however, many of these mutants were suppressed by *ndc80^ΔN-tail^* (Fig 6B and Fig S6B-C). These findings also raise the possibility that the N-terminus of Ndc80 associates directly with the interaction hub to regulate Mps1 localization.

**Figure 6.**
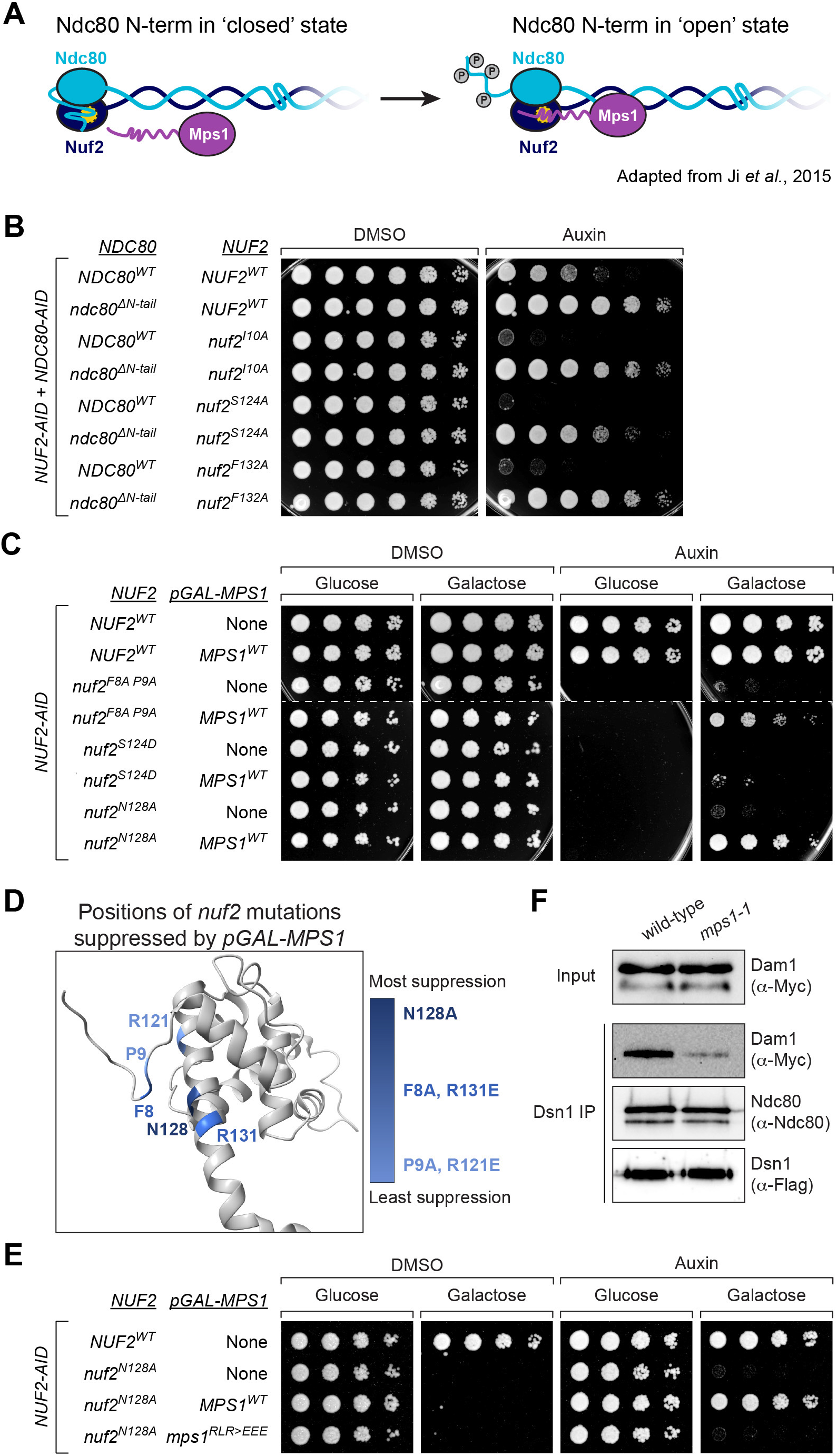
Increasing the level of Mps1 suppresses the lethality of *nuf2* mutants. (A) Model illustrating two states of the Ndc80 N-terminal tail: “closed”, in which the N-terminal tail of Ndc80 associates with Nuf2, precluding Mps1 binding, and “open”, in which the tail is altered, likely via phosphorylation, to allow Mps1-Nuf2 association^43^. (B) Yeast cell viability assay comparing *nuf2* mutant phenotypes in *NDC80^WT^* and *ndc80^ΔN-tail^* strains. Each is a “double AID” strain, containing both *NUF2-AID* and *NDC80-AID* alleles, as well as ectopic copies of *NUF2-3HA* and *NDC80-3HA* (*NUF2^WT^ NDC80^WT^*, M2998; *NUF2^WT^ ndc80^ΔN-tail^*, M2999; *nuf2^I10A^ NDC80^WT^*, M2654; *nuf2^I10A^ ndc80^ΔN-tail^*, M2755; *nuf2^S124A^ NDC80^WT^*, M2641; *nuf2^S124A^ ndc80^ΔN-tail^*, M2742; *nuf2^F132A^ NDC80^WT^*, M2841; *nuf2^F132A^ ndc80^ΔN-tail^*, M2774). Cells were serially diluted five-fold and spotted onto plates containing DMSO or 250 µM auxin. (C) Yeast cell viability assay to assess the effect of *pGAL*-driven overexpression of Mps1 in *nuf2* mutants. Strains contain *NUF2-AID* and ectopic copies of *NUF2-3HA*, without or with an ectopic *pGAL-MPS1* single integration vector allele, respectively (*NUF2^WT^*, M2038 and M3586; *nuf2^F8A^ ^P9A^*, M2042 and M3587; *nuf2^S124D^*, M2041 and M4619; *nuf2^N128A^*, M2414 and M3588). Cells were serially diluted five-fold and spotted onto plates containing either glucose (control) or galactose (induce over-expression of *MPS1*), as well as DMSO or 250 µM auxin. (D) Structure of *S. cerevisiae* Nuf2 (5TCS) illustrating the residues with mutants that are suppressed by expression of *pGAL-MPS1*. Darker shades of blue indicate increasing strength of the suppression. (E) Yeast cell viability assay, comparing *pGAL-MPS1^WT^* and *pGAL-mps1^RLR>EEE^* suppression of *nuf2^N128A^*. Strains contain *NUF2-AID* and an ectopic copy of *NUF2-3HA* (*NUF2^WT^*, M2038; *nuf2^N128A^*, M2414) or also contain a single, ectopic *pGAL-MPS1* allele (*nuf2^N128A^*with *MPS1^WT^*, M3588; *nuf2^N128A^* with *mps1^RLR>EEE^*, M4946). Cells were serially diluted five-fold and spotted onto plates containing either glucose or galactose, as well as DMSO or 250 µM auxin. (F) Detection of Dam1-9Myc association with the kinetochore via immunoprecipitation. Exponentially growing wild-type (M457) and *mps1-1* (M469) cultures expressing *DSN1-6His-3Flag*, *DAM1-9Myc* and *NDC80-3HA* were shifted to 37°C for 2 hours prior to harvest. Kinetochores were purified from lysates by anti-Flag immunoprecipitation (IP) and analyzed by immunoblotting.

Another way to modulate Mps1 levels at the kinetochore is through overexpression. Compellingly, overexpression of Mps1 rescues the cell viability defects of both *nuf2^F8A^ ^P9A^*and *nuf2^N128A^*, as well as another nearby mutation (Fig 6C & Fig S6D-E). In fact, when we examine the *nuf2* mutant alleles whose cell viability defects are suppressed by Mps1 overexpression, we find that the mutated residues spatially cluster on a surface of Nuf2 (Fig 6D). However, viability of *nuf2^S124D^*, the most penetrant single substitution *nuf2* mutant we tested, was not suppressed by Mps1 overexpression (Fig 6C) even though it is close in spatial proximity to F8, P9 and N128 (Fig 1A & Fig 6D). We suggest the increased severity of *nuf2^S124D^* may be due to physical displacement of the N-term loop of Nuf2 from its G-helix, leading to a complete disruption of the Mps1 binding site. The finding that the overexpression of Mps1 can suppress the detrimental effects of mutations in this portion of the Ndc80 complex implies that these mutations result in lethality due to reduced association between Mps1 and the Ndc80 complex. However, it was plausible that the rescue of these mutants by Mps1 overexpression was solely due to the restoration of spindle assembly checkpoint function^60–62^. This does not appear to be the case, as Mps1 overexpression still suppresses *nuf2* mutants even when the SAC is inactivated by deleting *MAD3* (Fig S6F). Furthermore, this suppression appears specific to Mps1 as overexpression of Ipl1 does not rescue the cell viability of these *nuf2* mutants (Fig S6G). Finally, the ability of Mps1 overexpression to suppress *nuf2* mutant growth defects appeared to be dependent on the ability of Mps1 to interact with the Ndc80 complex. While the overexpression of *MPS1^WT^* effectively suppresses the cell viability defects of the *nuf2* mutants, we found that the overexpression of *mps1^RLR>EEE^* or other mutations at the Ndc80 complex interface fail to achieve the same outcome (Fig 6E and Fig S6H-I). These findings indicate that overexpression of Mps1 alone is insufficient to suppress the *nuf2* mutants, but instead requires Mps1’s interaction with the Ndc80 complex. Moreover, the observation that certain *nuf2* mutants (e.g. *nuf2^F8A^ ^P9A^* and *nuf2^N128A^*) are susceptible to suppression by Mps1 overexpression suggests that these mutants disrupt, but do not entirely eliminate, the ability of Mps1 to bind the Ndc80 complex.

While modulating the levels of Mps1 at the kinetochore appears to suppress the *nuf2* CH domain mutants, how do we reconcile the observation that these mutants also display reduced Dam1 complex recruitment (as described in Fig 5A)? A potential explanation is that the localization of the Dam1 complex to the kinetochore is regulated by Mps1 activity, also suggested by previous work^31,36,63,64^. To investigate this hypothesis, we used a kinetochore co-IP assay^56^ in which kinetochores were immunoprecipitated, and co-purification of the Dam1 complex was determined by western blot. In line with the notion above, kinetochores purified from *mps1-1* cells lacking Mps1^53^ exhibited reduced association with the Dam1 complex (Fig 6F). These findings potentially elucidate how modulating Mps1 levels at the kinetochore can mitigate the effects caused by *nuf2* CH domain mutants, which disrupt the interaction hub and presumably lead to the loss of binding of multiple factors. Additionally, these results suggest an active role of Mps1 in recruiting a competing factor, the Dam1 complex, vying for the same kinetochore binding site.

### Mps1 is the critical factor recruited to the Ndc80 complex interaction hub

Finally, while modulating Mps1 levels appears to suppress the cell viability defects of the *nuf2* CH domain mutants, we wanted to more directly ask if the phenotypes of *nuf2* mutants are due specifically to diminished Mps1-kinetochore association. Here, we asked if restoring Mps1 localization, using rapamycin induced dimerization of FRB and FKBP12, would rescue the *nuf2* mutant phenotypes (Fig 7A). We generated strains containing *NUF2-AID*, *NDC80-FKBP12* and *MPS1-FRB* at their endogenous loci, and either *NUF2^WT^* or *nuf2^N128A^*expressed from an ectopic locus. Since tethering Mps1-FRB to the C-terminus of Ndc80 constitutively activates the SAC, causing cell inviability^65^, these strains also included a deletion of *MAD3*. Compellingly, the tethering of Mps1 to Ndc80 effectively restored the viability of *nuf2^N128A^* mutants (Fig 7B). These data argue that Mps1 needs to be bound to the Ndc80 complex to carry out its kinetochore biorientation function. In summary, our findings establish a pivotal “interaction hub” within the Ndc80 complex that orchestrates SAC signaling and microtubule attachment. Moreover, it underscores the indispensable role played by the kinetochore-associated pool of Mps1 in ensuring precise chromosome segregation.

**Figure 7.**
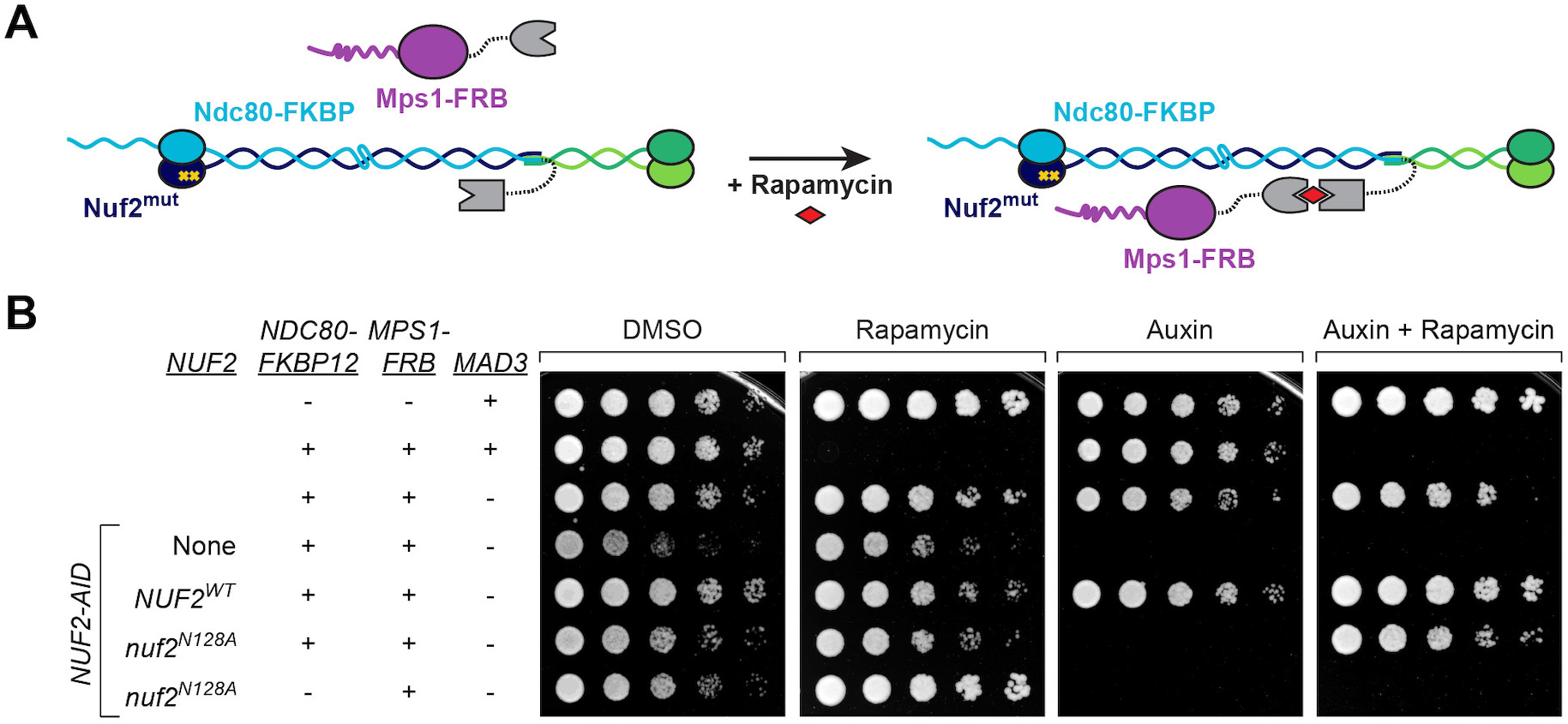
Tethering Mps1 to the kinetochore is sufficient to restore *nuf2* mutant viability. (A) Schematic of system to tether Mps1 to the kinetochore. In the presence of rapamycin, Mps1-FRB associates with Ndc80-FKBP12, artificially localizing Mps1 to the Ndc80 complex. (B) Yeast cell viability assay using the rapamycin-induced tethering system. *TOR1-1 fpr1Δ* (alone, M1375), followed by strains that also contain *MPS1-FRB* and *NDC80-FKBP12* tethering alleles *(MAD3*, M1463; *mad3Δ*, M4116) and *NUF2-AID*, with no *nuf2* covering allele (“None”, M4841) or with ectopic copies of *NUF2-3HA* (*NUF2^WT^*, M4908; *nuf2^N128A^*, M4912), and finally a control *nuf2^N128A^* strain lacking the *NDC80-FKBP12* allele (M4955). Cells were serially diluted five-fold and spotted onto plates containing DMSO, 50 ng/mL rapamycin, 250 µM auxin or 250 µM auxin plus 50 ng/mL rapamycin.

## DISCUSSION

In this work, we set out to examine the role that Nuf2’s CH domain plays in the fidelity of chromosome segregation. Through extensive mutational analysis, we pinpointed a conserved “interaction hub” within Nuf2’s CH domain, formed by portions of its N-terminal loop and G-helix. Importantly, this patch serves as the binding site for Mps1 within the yeast Ndc80 complex, but also associates with numerous other factors. Consequently, mutants disrupting this hub exhibit defects in spindle assembly checkpoint function and display severe errors in chromosome segregation. The array of interactions occurring at this specific site on the Ndc80 complex appears critical for the functional regulation of Mps1’s association with the kinetochore. In fact, restoring Mps1-Ndc80 complex association completely rescues the cellular defects associated with mutations at this site. Overall, this work sheds light on mechanisms cells use to regulate Mps1 at the kinetochore, and also underscores the essential role of Mps1 in kinetochore biorientation and precise chromosome segregation.

### Mps1 is the critical factor that associates with the Ndc80 complex’s interaction hub

Our work describes a previously uncharacterized region of Nuf2’s CH domain that is responsible for interacting with segments of various proteins. This list encompasses the N-terminus of the kinase Mps1, the C-terminus of the Dam1 protein, potentially the N-terminal tail of the Ndc80 protein, and the N-terminus of the kinase Ipl1^19,20,42,43^ (this work as well as concurrent work by Zahm *et al*, Pleuger *et al*; personal communication S. Harrison and S. Westermann), among potentially other yet-to-be-identified factors. Notably, mutations affecting this interaction hub lead to cell inviability and substantial errors in kinetochore biorientation and chromosome segregation. Interestingly, our findings suggest that these severe phenotypes do not result from disrupting the binding of multiple factors to this site. Instead, our observation that specifically restoring Mps1-Ndc80 complex association, whether through overexpression or tethering via FRB-FKBP, implies that Mps1 might be the sole essential factor binding to this interaction hub. It remains possible that Mps1’s role may involve, in part, recruiting and/or regulating these additional factors when bound to the Ndc80 complex. Our observation that Mps1 appears to recruit the Dam1 complex aligns with this latter concept, and is consistent with previous work^31,36,63,64^. In any case, our identification of the Mps1 binding site on the Ndc80 complex highlights the crucial role that kinetochore-bound Mps1 plays in error correction, in addition to its well-studied function in spindle assembly checkpoint activation. This error correction function of Mps1 is essential for cell viability, revealing a more significant role than we previously recognized.

### The activity of Mps1 at the kinetochore is subject to regulation at multiple levels

We propose the following working model, building upon the excellent work of others (reviewed in ^3,66^), outlining the steps from the initial microtubule attachment to biorientation (Fig 8). Prior to initial attachment to spindle microtubules, the Ndc80 complex is in a ‘closed’ conformation with the N-terminal tail of Ndc80 folded back and occluding the association of other factors to this interaction hub^43^. Both error correction kinases, Ipl1 and Mps1, can phosphorylate the N-terminal tail^7,11–13,27–31^, which changes Ndc80 to an ‘open’ conformation^43^, allowing binding of Mps1. Kinetochore-associated Mps1 can now carry out its well-described function in spindle assembly checkpoint activation^33–35,37,44,45^. Importantly, this pool of Mps1 also appears essential for achieving kinetochore biorientation and ensuring accurate chromosome segregation. This is likely achieved through a tension-sensitive error correction mechanism^31–37^. However, the precise working mechanism of this process remains unclear (further discussion below). In part, this error correction function of Mps1 may involve recruiting the Dam1 complex^31,36,63,64^. Regardless, once attachments are correctly bioriented and no longer susceptible to error correction-mediated destabilization, the Dam1 complex establishes stable association with the Ndc80 complex, thereby displacing Mps1 from its binding site (Zahm *et al.*, Pleuger *et al.*; personal communication S. Harrison and S. Westermann). In this manner, Mps1 seems to facilitate the recruitment of its own competitive inhibitor. Another mechanism through which Mps1 regulates its kinetochore localization is autophosphorylation^58,67^. Furthermore, competition between Mps1 and microtubules for binding to the Ndc80 complex is proposed as an additional means to displace Mps1^42,43^, although Mps1 binding does not appear to be mutually exclusive to microtubule binding^67^. These cumulative events subsequently deactivate spindle assembly checkpoint signaling and also likely terminate Mps1’s participation in error correction. While this model provides insights into certain aspects of this process, several crucial questions remain.

**Figure 8.**
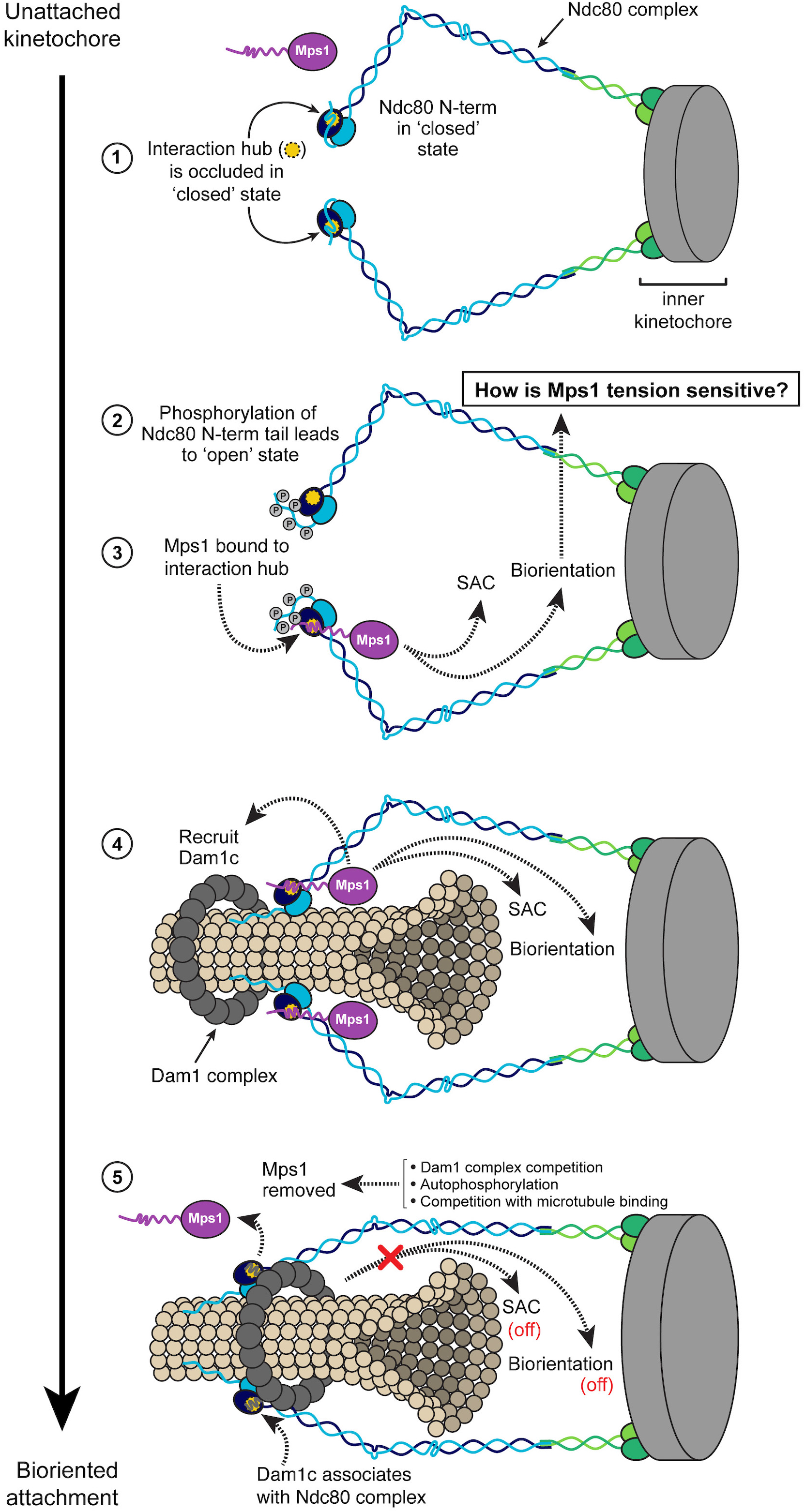
Model of the transition from unattached kinetochore to bioriented attachment, incorporating known protein-protein associations and events that may occur at the interaction hub on Nuf2. The interaction hub on Nuf2 (yellow) interacts with Mps1 (purple) and the Dam1 complex (dark grey), and potentially with the N-terminal tail of Ndc80 (light blue). The microtubule is shown in tan. 1. In the unattached kinetochore state, the N-terminus of Ndc80 folds over Nuf2 to occlude the interaction hub (“closed state”), precluding association of Mps1 with the kinetochore. 2. Movement of the Ndc80 N-terminal tail away from the Nuf2 interaction hub (possibly via phosphorylation by Ipl1 or Mps1) leads to the “open state”, allowing access to other factors. 3. Mps1 associates with the Ndc80 complex at the Nuf2 hub, triggering the SAC and participating in sister chromatid biorientation through an as yet undetermined mechanism that is likely tension sensitive. 4. Mps1 also promotes association of the Dam1 complex with Ndc80 complex, which ultimately allows for stable kinetochore-microtubule interaction and the formation of bioriented attachments. 5. Once the bioriented state is achieved, Mps1 is removed from the kinetochore via competition by the Dam1 complex for the interaction hub, autophosphorylation^67^ and/or competition with microtubule binding^42,43^. Loss of Mps1 at the kinetochore also turns off the SAC.

One key question is how does Mps1’s activity at kinetochores contribute to biorientation? Additionally, what mechanisms regulate these activities in response to tension, a critical signal that cells use to discern correct from incorrect attachments? It has been well-demonstrated that Mps1 phosphorylates sites on the N-terminal tail of Ndc80, leading to attachment destabilization. Notably, Mps1’s phosphorylation of Ndc80 is stimulated on kinetochores lacking tension^31^. However, disrupting the known Mps1 phosphorylation sites, even in conjunction with a spindle assembly checkpoint mutant, does not substantially impact cell viability^30,31^. Furthermore, the N-terminal tail of the Ndc80 protein is dispensable for viability in yeast^30,68,69^. These observations are in contrast to the *nuf2* and *mps1* mutants described here, which exhibit complete inviability. This discrepancy suggests that additional mechanisms must be involved in this process. Mps1 has been implicated in two other pathways that promote biorientation: the activation of the kinase Bub1^32,70^ and the recruitment of pericentromeric localization of Sgo1^70–75^. The interplay between these pathways, particularly in the context of our new mutants disrupting kinetochore-associated Mps1, requires further investigation. It’s worth noting that Bub1 and Sgo1 are non-essential proteins, and furthermore, the role of tension in these processes remains unclear.

Several key questions surround the regulation of the kinetochore-bound Mps1 pathway by tension. Given the proximity of Mps1 to the microtubule binding domains of the kinetochore, spatial separation of substrates from Mps1, as proposed for Aurora B^76^, seems unlikely. Instead, tension-dependent changes in substrate conformation, as recently demonstrated for Ipl1^77^, provide a plausible mechanism by which tension suppresses phosphorylation of critical targets. Alternatively, Mps1’s kinase activity or localization could be influenced by a kinetochore factor, potentially a portion of the Ndc80 complex, and this regulation may be responsive to tension. For instance, the displacement of Nuf2’s N-terminal loop from its G-helix (similar to the *nuf2^S124D^* mutant) either via tension or another mechanism appears to be an intriguing way that cells could modulate Mps1 activity. This notion gains support from the observation that the assembly of the outer kinetochore is essential for generating most, if not all, cellular Mps1 activity^55^. Additionally, the interplay between Aurora B and Mps1 activities remains an open question.

In summary, our work identifies a critical “interaction hub” that governs essential kinetochore functions of the kinase Mps1. These functions encompass its broadly studied role in spindle assembly checkpoint signaling, but, notably, emphasize the significance of Mps1 in establishing kinetochore biorientation. Our findings clarify the mechanisms that govern kinetochore-associated Mps1 and how these mechanisms contribute to the fidelity of chromosome segregation.

## ACKNOWLEDGEMENTS

The authors declare no competing financial interests. We are grateful to Stefan Westermann, Richard Pleuger, Steve Harrison and Jake Zahm for sharing data prior to publication. We thank Sue Biggins, Leon Chan, Trisha Davis, Adèle Marston, Michael Stewart, Andrew Murray, and Frank Uhlmann for providing plasmids and/or strains and Arshad Desai for providing antibodies. In addition, we would like to thank members of the Miller lab, Stefan Westermann and Steve Harrison for helpful discussions and critical reading of the manuscript. This work was supported by 5 For the Fight (to M.P.M), Pew Biomedical Scholars (to M.P.M), and NIH grant R35GM142749 (to M.P.M).

## METHODS

### Yeast strains and media

All yeast strains used are listed in Table S1 and are isogenic in the W303 background^78^. Standard media and microbial techniques^79^ were used, as well as standard yeast genetic methods for strain construction^80,81^.

Single integration vectors (SIV) pM93 (*HIS3*), pM94 (*URA3*), pM95 (*TRP1*), and pM96 (*LEU2*) were provided by Leon Chan and were used in the construction of the *NUF2*, *NDC80*, *MPS1* and *IPL1* plasmids described below. The genomic *NUF2-AID* allele *NUF2-3V5-IAA7:KanMX* was constructed using pM66 (*3V5-IAA7:KanMX* plasmid; Leon Chan) as a template, and subsequent conversion to *NUF2-3V5-IAA7:NATMX* was performed as in ^82^. A single integration *LEU2* vector containing *pNUF2-NUF2-3HA* (pM604) was constructed by first tagging endogenous *NUF2* with 3HA from pM9 (*3HA:HIS3MX6*)^83^, then using the *pNUF2-NUF2-3HA* strain as a template for plasmid construction. Mutants of *NUF2* were constructed by mutagenizing pM604 as described in ^84,85^. The *NDC80-3V5-IAA7:KanMX* is described in ^39^. A *TRP1* single integration vector containing *pNDC80-NDC80-3HA* (pM270) was constructed in a similar manner to *NUF2-3HA* above, and *pNDC80-ndc80^ΔN-tail^-3HA* (pM1342) was subsequently constructed using pM270 as a template for mutagenesis as in ^84,85^.

The *MPS1-3V5:KanMX* allele was constructed using pM32 (*V5:KanMX* tagging vector; Leon Chan) as a template, and was marker converted to *MPS1-3V5:HphMX* as in ^82^. *pGAL10-MPS1* and *pGAL10-IPl1 HIS3* single integration vectors were constructed by genomic PCR amplification of the respective ORFs, followed by Gibson assembly into pM93, downstream of the *pGAL10* promoter^86^. The *pGAL10* promoter was replaced with genomic PCR-amplified native *MPS1* promoter to construct pM1595 (*pMPS1-MPS1 HIS3* SIV), and subsequent cloning produced pM1718 (*pMPS1-MPS1 URA3* SIV). Mutants of *MPS1* were constructed in these plasmids by standard cloning or by mutagenizing pM1489, pM1595 and pM1718 as described in ^84,85^. All plasmids and primers are listed in Table S2. Further details regarding plasmid and strain construction are available upon request.

Integration plasmids containing *pGPD1-TIR1* (pM74 for integration at *LEU2*, pM76 for *HIS3*, and pM78 for *TRP1*) were provided by Leon Chan; the *URA3* version was constructed from pM74 and the pM94 *URA3* single integration vector. *SPC110-mCherry:HphMX* and *ASK1-YFP:HIS3* were provided by Trisha Davis, and *BUB1-GFP:KanMX*, *MTW1-mCherry:HphMX* by Sue Biggins. Construction of *pCUP1-GFP-LacI* is described in ^87^, *CEN III::lacO:TRP1* is described in ^88^, and *CEN IV::lacO:TRP1* was provided by Andrew Murray. Strains containing the previously described *pMET-CDC20* allele were provided by Frank Uhlmann^89^, and the *CDC20-AID* allele by Adele Marston. *mps1-1* and *dad1-1* temperature sensitive alleles were provided by Andrew Murray and Sue Biggins, respectively. The *pGAL-MPS1-Myc:URA3* and *mad2Δ* alleles were provided by Andrew Murray, *mad3Δ* is described in ^90^, and *DSN1-6His-3Flag* in ^56^. *TOR1-1*, *fpr1Δ*, *MPS1-FRB:KanMX*, and *NDC80-FKBP12:HISMX* are described in ^65,91^. The Dam1-9Myc allele was provided by Sue Biggins.

### Auxin inducible degron

The auxin inducible degron (AID) system was used as described in ^92^. Cells expressed C-terminal fusions of *NUF2*, *NDC80*, or *CDC20* to the gene for an auxin responsive protein (*IAA7*) at each respective endogenous locus. Cells also expressed *TIR1*, which is required for auxin-induced degradation. For cell viability assays, 250 µM auxin (indole-3-acetic acid; Sigma Aldrich) dissolved in DMSO was top-plated on agar to induce degradation of the AID-tagged protein. Cells for microscopy were treated with 500 µM auxin in liquid media at the specified time prior to fixation. Auxin was added to liquid media 2 hours prior to harvesting cells for immunoprecipitation analysis.

### Spot dilution yeast viability assays

Desired strains were grown overnight for two days on YPA plus 2% dextrose plates (YPAD). Equal amounts of each strain were then used for a 1:5 dilution series and spotted onto YPAD + DMSO, YPAD + Auxin, YPAD + Auxin and 6.5 µg/mL benomyl, YPA Gal (YPA + 2% galactose) + DMSO, or YPA Gal + Auxin. Plates were incubated at 23°C for 2 to 4 days, unless otherwise indicated for temperature sensitive alleles (30°C or 37°C, as noted in figure legends).

### Cell fixation, imaging conditions and image analysis

Conditions for growing cells for each type of imaging are described in brief in the figure legends. For most experiments, cells were grown to log phase in standard YPAD liquid medium at room temperature (approximately 20°C). Where indicated, cells were treated with auxin (500 µM) or auxin and nocodazole (10 µM). In some cases, an α-factor arrest was used to synchronize cells (1 µg/mL for 3 hours at room temperature; using *bar1-1* strains), followed by three washes in YPAD + 2% DMSO and resuspension in fresh YPAD media for the release. For the biorientation assay, cells were arrested with α-factor in synthetic liquid medium lacking methionine, then released into the *pMET-CDC20* arrest in the presence of methionine at 30°C. 8 mM methionine was subsequently added every thirty minutes until harvest.

Fixation was performed in 3.7% formaldehyde in 100 mM phosphate buffer (pH 6.4) for 5 minutes. Cells were washed once with 100 mM phosphate (pH 6.4) and resuspended in 100 mM phosphate, 1.2 M sorbitol buffer (pH 7.5) and permeabilized with 1% Triton X-100 stained with 1 µg/mL DAPI (4’, 6-diamidino2-phenylindole; Molecular Probes). Cells were imaged with a DeltaVision Ultra microscope with a 60X objective (NA = 1.42), equipped with a sCMOS digital camera. Fifteen Z-stacks (0.3 micron apart) were acquired, and frames were deconvolved using standard settings. Image stacks were maximally projected. softWoRx image processing software was used for image acquisition and processing.

Quantitation of microscopy images was performed as indicated in the figure legends. For the Bub1-GFP assay for SAC activity, only mononucleate cells with a single Mtw1-mCherry signal were counted (indicating collapse of the spindle from nocodazole treatment). Note that *nuf2* mutants often increased the number of cells with two Mtw1 signals, despite nocodazole treatment, and also displayed increased bypass of the *CDC20-AID* arrest (Fig S3A-C). Collectively, this reduced the number of cells in each field of view available for scoring the presence of Bub1-GFP at the kinetochore relative to *NUF2^WT^*. Numbers of cells counted are indicated in the figure legend. For the biorientation assay in Fig 3B, only mononucleate cells in metaphase were counted. For chromosome segregation assays in Fig 3C and Fig 4D, only binucleate cells that had undergone anaphase (approximate spindle length ≥ 4 µM) were counted.

Spindle length in Fig S3A was calculated by drawing a line in FIJI (Image J, NIH) that connected the edges of the Spc110-mCherry signals from the two poles. Spindle pole body separation in Fig S4B was calculated by the percentage of cells that displayed two Spc110-mCherry signals at 90 minutes following the temperature shift. Ask1-YFP intensities in Fig 5A were calculated by drawing a line in FIJI through the Spc110 spindle pole signals and performing plot profile analysis of the YFP signal across this line. The maximum peak intensity proximal to each pole was then determined (two signals for each cell, which were sometimes asymmetric, especially in the *nuf2* mutant). Data points in the right graph in Fig 5 represent individual Ask1-YFP peak intensities, two for each cell.

### Kinetochore immunoprecipitations

Native kinetochores were purified from asynchronously, exponentially growing *S. cerevisiae* cells containing Dsn1-6His-3Flag by immunoprecipitation with α-Flag, essentially as described in ^56^. Cells were grown in standard YPAD medium. For strains containing *NUF2-AID*, cells were treated with 500 μM auxin 2 hours prior to harvest. For examining kinetochore enrichment of Dam1-9Myc in WT and *mps1-1* strains, cells were shifted to 37°C for 2 hours prior to harvest. Protein lysates were prepared by mechanical disruption in the presence of lysis buffer using glass beads and a beadbeater (Biospec Products). Lysed cells were resuspended in buffer H (BH) (25 mM HEPES pH 8.0, 2 mM MgCl2, 0.1 mM EDTA, 0.5 mM EGTA, 0.1% NP-40, 15% glycerol with 150 mM KCl containing protease inhibitors (at 20 μg/mL final concentration for each of leupeptin, pepstatin A, chymostatin and 200 μM phenylmethylsulfonyl fluoride) and phosphatase inhibitors (0.1 mM Na-orthovanadate, 0.2 μM microcystin, 2 mM β-glycerophosphate, 1 mM Na pyrophosphate, 5 mM NaF) followed by centrifugation at 16,100 g for 30 min at 4°C to clarify the lysate. Dynabeads conjugated with α-Flag antibodies were incubated with extract for 3 hours with constant rotation, followed by three washes with BH containing protease inhibitors, phosphatase inhibitors, 2 mM dithiothreitol (DTT) and 150 mM KCl. Beads were further washed twice with BH containing 150 mM KCl and protease inhibitors. Associated proteins were eluted from the beads by boiling in 2x SDS sample buffer.

### Immunoblotting

For immunoblot analysis in Fig S1C, cell lysates were prepared by mechanical disruption with glass beads in 2x SDS-PAGE sample buffer (0.4mM EDTA, 4% SDS, 125 mM Tris pH6.8, 20% glycerol, 2% bromophenol blue). Lysates for immunoprecipitation were described above. Standard procedures for sodium dodecyl sulfate-polyacrylamide gel electrophoresis (SDS-PAGE) and immunoblotting were followed as described in ^93,94^. A nitrocellulose membrane (Bio-Rad) was used to transfer proteins from polyacrylamide gels. Commercial monoclonal antibodies used for immunoblotting were as follows: a-HA, 12CA5 (Roche, 1:2000), a-Flag, M2 (Sigma Aldrich, 1:3000), a-V5 (Invitrogen; 1:5000), a-Pgk1 (Abcam; 1:5000). The Ndc80 polyclonal antibody was a gift from Arshad Desai and was used at 1:10,000. Secondary antibodies used were a sheep anti-mouse and goat anti-rabbit conjugated to horseradish peroxidase (HRP, GE Biosciences; 1:10,000 dilution). Antibodies were detected using the SuperSignal West Dura Chemiluminescent Substrate (Thermo Scientific).

### Statistics

GraphPad Prism version 10 was used for statistical analysis. Data normality was assumed for all experiments. Student’s *t* test was used for comparisons between strains, as indicated in the figure legends. In the text, mean ± standard deviation are reported.

### Multiple sequence alignments

Fungal Nuf2 proteins were identified using a PSI-BLAST^95^ search on NCBI. Multiple sequence alignments of the entire proteins were generated with ClustalOmega default parameters and displayed in JalView 2^96^. Species are listed in the figures and/or legends. Conservation scores shown on structures were generated in ChimeraX 1.5, using an entropy-based measure from AL2CO^97^.

