## Supplemental Files for "An interaction hub on Ndc80 complex facilitates dynamic recruitment of Mps1 to yeast kinetochores to promote accurate chromosome segregation"

**SUPPLEMENTATAL INFORMATION**

Figures S1-S6 and Tables S1, S2.

#### Figure S1

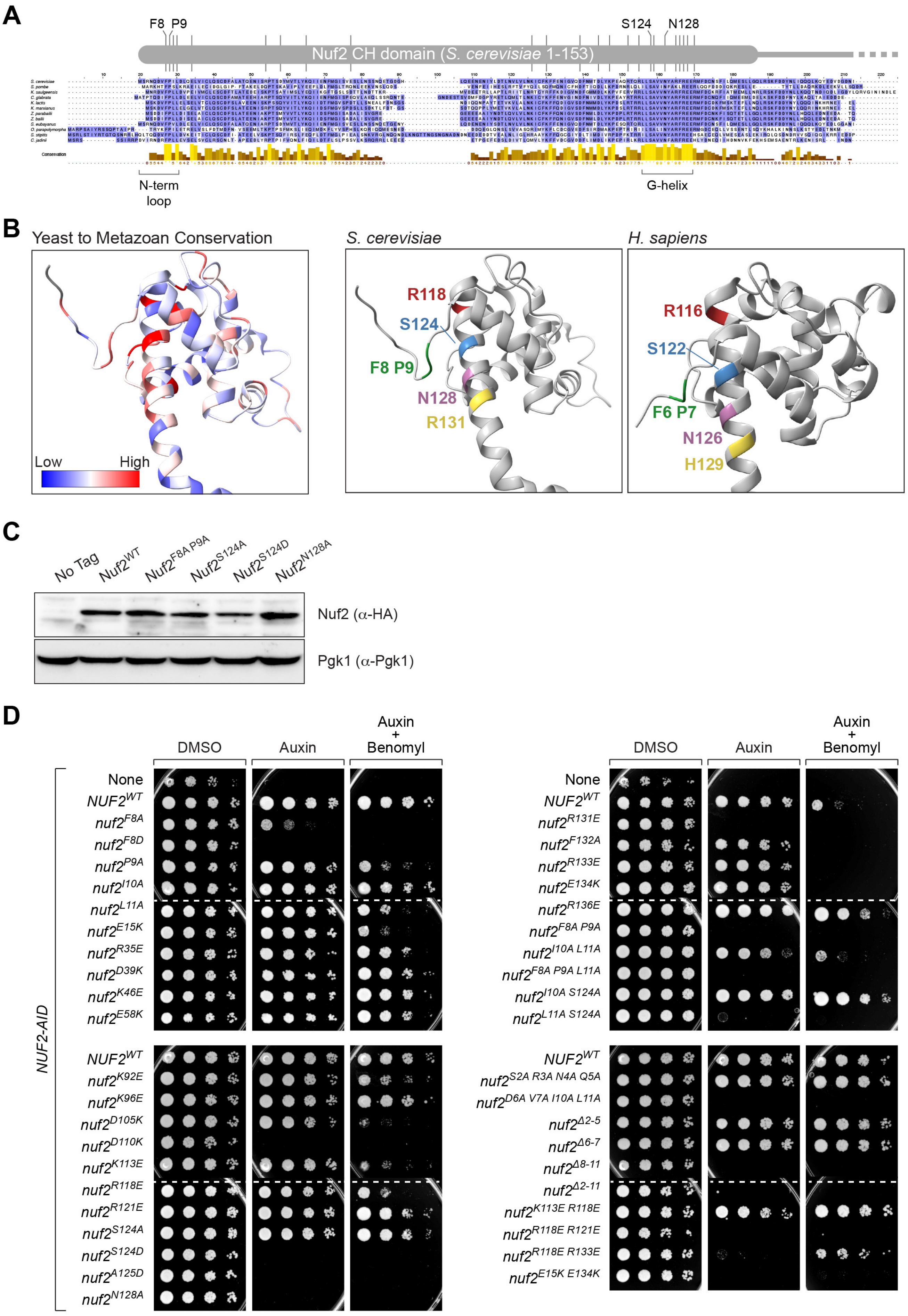

A
F8 P9
S124 N128
Nuf2 CH domain (S. cerevisiae 1-153)
Conservation
N-term loop
G-helix
B
Yeast to Metazoan Conservation
Low
High
S. cerevisiae
R118
S124
F8 P9
N128
R131
H. sapiens
R116
S122
F6 P7
N126
H129
C
No Tag
Nuf2WT
Nuf2F8A P9A
Nuf2S124A
Nuf2S124D
Nuf2N128A
Nuf2 (α-HA)
Pgk1 (α-Pgk1)
D
DMSO
Auxin
Auxin + Benomyl
NUF2-AID
None
NUF2WT
nuf2F8A
nuf2F8D
nuf2P9A
nuf2I10A
nuf2L11A
nuf2E15K
nuf2R35E
nuf2D39K
nuf2K46E
nuf2E58K
NUF2WT
nuf2K92E
nuf2K96E
nuf2D105K
nuf2D110K
nuf2K113E
nuf2R118E
nuf2R121E
nuf2S124A
nuf2S124D
nuf2A125D
nuf2N128A
None
NUF2WT
nuf2R131E
nuf2F132A
nuf2R133E
nuf2E134K
nuf2R136E
nuf2F8A P9A
nuf2I10A L11A
nuf2F8A P9A L11A
nuf2I10A S124A
nuf2L11A S124A
NUF2WT
nuf2S2A R3A N4A Q5A
nuf2D6A V7A I10A L11A
nuf2Δ2-5
nuf2Δ6-7
nuf2Δ8-11
nuf2Δ2-11
nuf2K113E R118E
nuf2R118E R121E
nuf2R118E R133E
nuf2E15K E134K

**Figure S1. Conservation of the Nuf2 CH domain and phenotypes of *nuf2* mutants.**

(A) Multiple sequence alignment showing conservation of the Nuf2 CH domain (residues 1-153) from fungal species, colored by BLOSUM62 scores. Numbering corresponds to that of the *S. cerevisiae* protein. From top to bottom: *Saccharomyces cerevisiae*, *Schizosaccharomyces pombe*, *Kazachstania saulgeensis*, *Candida glabrata*, *Kluyveromyces lactis*, *Kluyveromyces marxianus*, *Zygosaccharomyces parabailii*, *Zygosaccharomyces bailii*, *Saccharomyces eubayanus*, *Ogataea parapolymorpha*, *Scheffersomyces stipitis*, *Cyberlindnera jadinii*. Cartoon on the top indicates positions of *nuf2* mutant alleles (vertical lines). Degree of conservation is shown in the histogram.

(B) Left: Structure of *S. cerevisiae* Nuf2 from [49] (5TCS), illustrating conservation from yeast to metazoans, viewed using ChimeraX, on a scale of -2.5 to 2.5. Species included: *Saccharomyces cerevisiae*, *Caenorhabditis elegans*, *Danio rerio*, *Xenopus laevis*, *Gallus gallus*, *Mus musculus*, *Rattus norvegicus*, *Homo sapiens.* Right: Comparison of Nuf2 structures from *S. cerevisiae* (5TCS) and *H. sapiens* (2VE7; Ciferri et al, 2008), with highly conserved residues highlighted.

(C) Structural integrity of Nuf2 mutant proteins. Exponentially growing *NUF2-AID* strains with ectopic *NUF2-3HA* (No covering allele, “No tag”, M1889; *NUF2*$^{WT}$, M2038; *nuf2*$^{F8A\ P9A}$, M2042; *nuf2*$^{S124A}$, M2040; *nuf2*$^{S124D}$, M2041, *nuf2*$^{N128A}$, M2414) were used to prepare lysates that were analyzed by immunoblotting. Pgk1 was used as a loading control.

(D) Yeast cell viability assay with *nuf2* mutant alleles. Strains carry *NUF2-AID* and an ectopic copy of *NUF2-3HA* (No covering allele, “None”, M1889; *NUF2*$^{WT}$, M2038; *nuf2*$^{F8A}$, M2151; *nuf2*$^{F8D}$, M3996; *nuf2*$^{P9A}$, M2152; *nuf2*$^{I10A}$, M2191; *nuf2*$^{L11A}$, M2192; *nuf2*$^{E15K}$, M3443; *nuf2*$^{R35E}$, M2227; *nuf2*$^{D39K}$, M3441; *nuf2*$^{K46E}$, M2228; *nuf2*$^{E58K}$, M3444; *nuf2*$^{K92E}$, M2229; *nuf2*$^{K96E}$, M2230; *nuf2*$^{D105K}$, M2413; *nuf2*$^{D110K}$, M2544; *nuf2*$^{K113E}$, M2153; *nuf2*$^{R118E}$, M2154; *nuf2*$^{R121E}$, M2200; *nuf2*$^{S124A}$, M2040; *nuf2*$^{S124D}$, M2041; *nuf2*$^{A125D}$, M2633; *nuf2*$^{N128A}$, M2414; *nuf2*$^{R131E}$, M2201; *nuf2*$^{F132A}$, M2545; *nuf2*$^{R133E}$, M2202; *nuf2*$^{E134K}$, M3442; *nuf2*$^{R136E}$, M2203; *nuf2*$^{F8A\ P9A}$, M2042; *nuf2*$^{I10A\ L11A}$, M3750; *nuf2*$^{F8A\ P9A\ L11A}$, M2262; *nuf2*$^{I10A\ S124A}$, M3751; *nuf2*$^{L11A\ S124A}$, M3752; *nuf2*$^{S2A\ R3A\ N4A\ Q5A}$, M2043; *nuf2*$^{D6A\ V7A\ I10A\ L11A}$, M2044; *nuf2*$^{\Delta 2\text{-}5}$, M2048; *nuf2*$^{\Delta 6\text{-}7}$, M2046; *nuf2*$^{\Delta 8\text{-}11}$, M2047; *nuf2*$^{\Delta 2\text{-}11}$, M2045; *nuf2*$^{K113E\ R118E}$, M2323; *nuf2*$^{R118E\ R121E}$, M2264; *nuf2*$^{R118E\ R133E}$, M2263; *nuf2*$^{E15K\ E134K}$, M3507). Cells were serially diluted five-fold and spotted onto plates containing DMSO, 250 µM auxin or 250 µM auxin + 6.5µg/mL benomyl.

### Figure S2

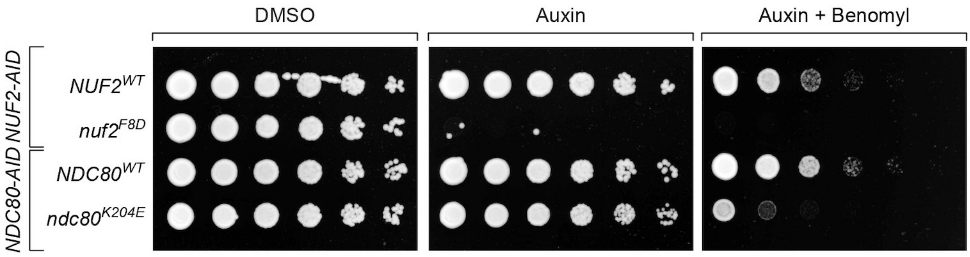
DMSO
Auxin
Auxin + Benomyl
NUF2-AID
NDC80-AID
$NUF2^{WT}$
$nuf2^{F8D}$
$NDC80^{WT}$
$ndc80^{K204E}$

**Figure S2. Comparison of phenotypes of a single negatively charged amino acid introduced into *NUF2* vs. *NDC80*.**

Yeast cell viability assay comparing single amino acid substitution *nuf2* and *ndc80* mutant alleles. Strains carry either a *NUF2-AID* or an *NDC80-AID* and an ectopic copy of *NUF2-3HA* or *NDC80-3HA* ($NUF2^{WT}$, M2038; $nuf2^{F8D}$, M3996; $NDC80^{WT}$, M692; $ndc80^{K204E}$, M693). Cells were serially diluted five-fold and spotted onto plates containing DMSO, 250 μM auxin or 250 μM auxin + 6.5 μg/mL benomyl. Note that a single amino acid substitution in Nuf2's N-term loop causes a dramatically larger growth defect than a similar mutation in Ndc80's CH domain believed to impair microtubule binding[7,10].

Figure S3

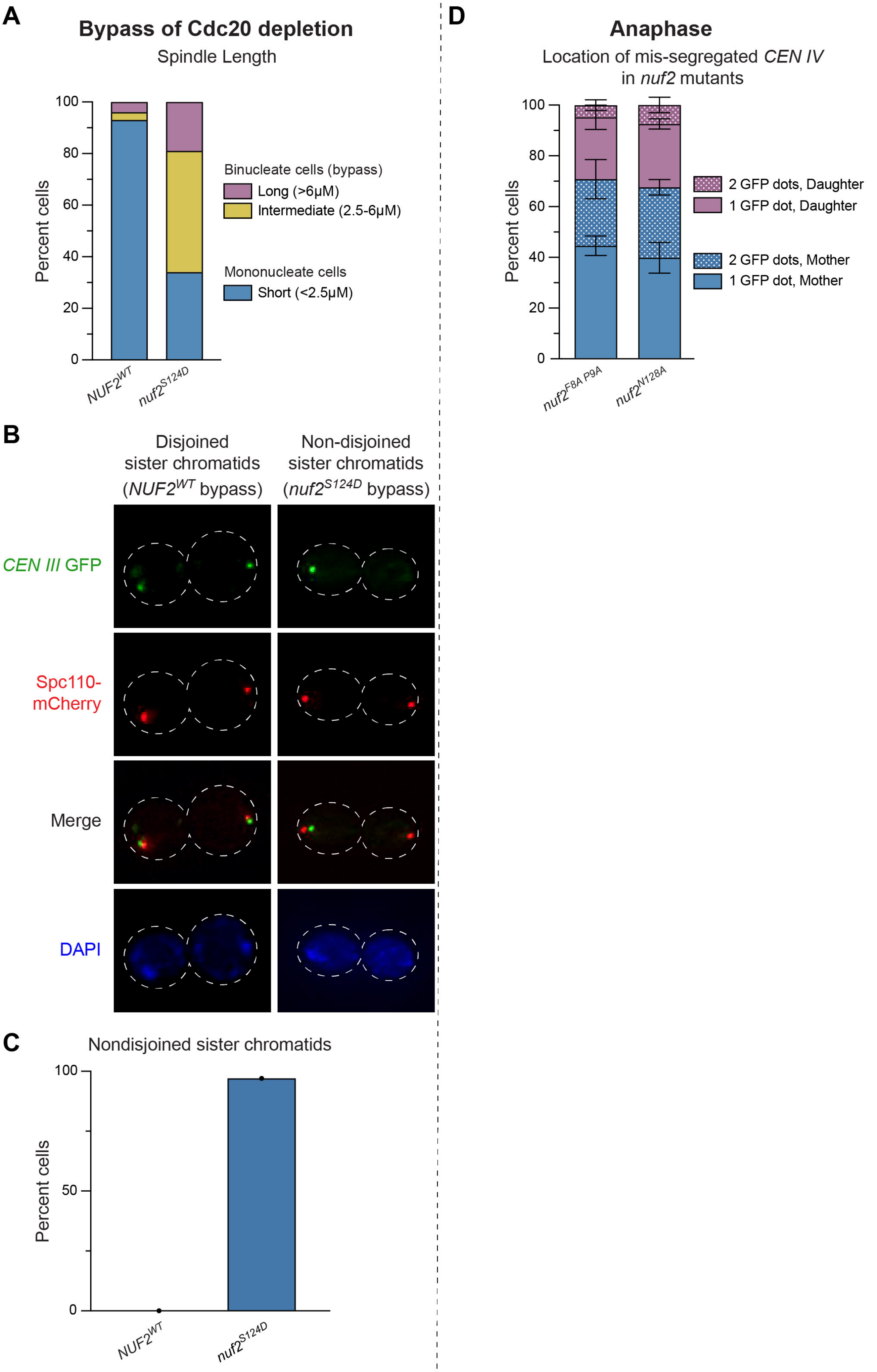

A
Bypass of Cdc20 depletion
Spindle Length
Percent cells
100
80
60
40
20
0
NUF2WT
nuf2S124D
Binucleate cells (bypass)
Long (>6μM)
Intermediate (2.5-6μM)
Mononucleate cells
Short (<2.5μM)
B
Disjoined sister chromatids (NUF2WT bypass)
Non-disjoined sister chromatids (nuf2S124D bypass)
CEN III GFP
Spc110-mCherry
Merge
DAPI
C
Nondisjoined sister chromatids
Percent cells
100
50
0
NUF2WT
nuf2S124D
D
Anaphase
Location of mis-segregated CEN IV in nuf2 mutants
Percent cells
100
80
60
40
20
0
nuf2F8A P9A
nuf2N128A
2 GFP dots, Daughter
1 GFP dot, Daughter
2 GFP dots, Mother
1 GFP dot, Mother

**Figure S3. Bypass of Cdc20 depletion is observed in *nuf2* mutants, as well as mis-localized *CEN IV* GFP in anaphase.**

*Bypass of Cdc20 depletion (left)*

(A) Bypass of a *pMET-CDC20* arrest by the *nuf2*$^{S124D}$ mutant. Strains contain *NUF2-AID* and ectopic *NUF2-3HA* (*NUF2*$^{WT}$, M3753; *nuf2*$^{S124D}$, M3757) as well as the *SPC110-mCherry* spindle pole body marker. Distance between mCherry signals was calculated as the spindle length. Graph shows the percent of cells in each of three categories: arrested, mononucleate cells with short spindles (<2.5uM), and "bypass" binucleate cells with intermediate (2.5-6uM) or long spindles (>6uM); n = 61-113 cells for each genotype. The bypass observed in these cells likely artificially or partially masks the true level of biorientation defect (Fig 3B). Cells with bioriented sister chromatids will remain arrested (and thus be counted), whereas cells that bypassed the arrest likely did so due to significant biorientation defects, but are not counted as part of the assay, since only mononucleate cells are included in the analysis.

(B) Micrograph examples showing position of sister chromatids in cells that bypassed a *CDC20-AID* arrest in strains carrying *CEN III* marked with GFP (*CEN III:lacO LacI-GFP*), *SPC110-mCherry* (spindle pole marker), *NUF2-AID* and ectopic *NUF2-3HA* (*NUF2*$^{WT}$, M3874; *nuf2*$^{S124D}$, M3878). In the *NUF2*$^{WT}$ strain, cells that bypass the arrest display disjoined sister chromatids; in the *nuf2*$^{S124D}$ mutant example, bypassed cells demonstrate non-disjunction of sister chromatids, accompanied by significant variations in nuclear masses. These findings support the notion that in *nuf2* mutants, nearly all sister chromatids are attached to one of the two spindle poles, and thereby lack any constraints on spindle elongation.

(C) Graph illustrating the percentage of cells from (B) that show non-disjoined sister chromatins following bypass of the *CDC20* arrest (n = 22-32 cells for each genotype).

*Anaphase (right)*

(D) Quantitation of the location of mis-localized *CEN IV* GFP from *NUF2*$^{F8A\ P9A}$ (M4475) vs. *nuf2*$^{N128A}$ (M4479), shown in Figure 3C. The percent cells with 1 vs. 2 *CEN IV* GFP signals in either the mother or the daughter is shown. Error bars indicate the standard deviation among 3 replicates.

Figure S4

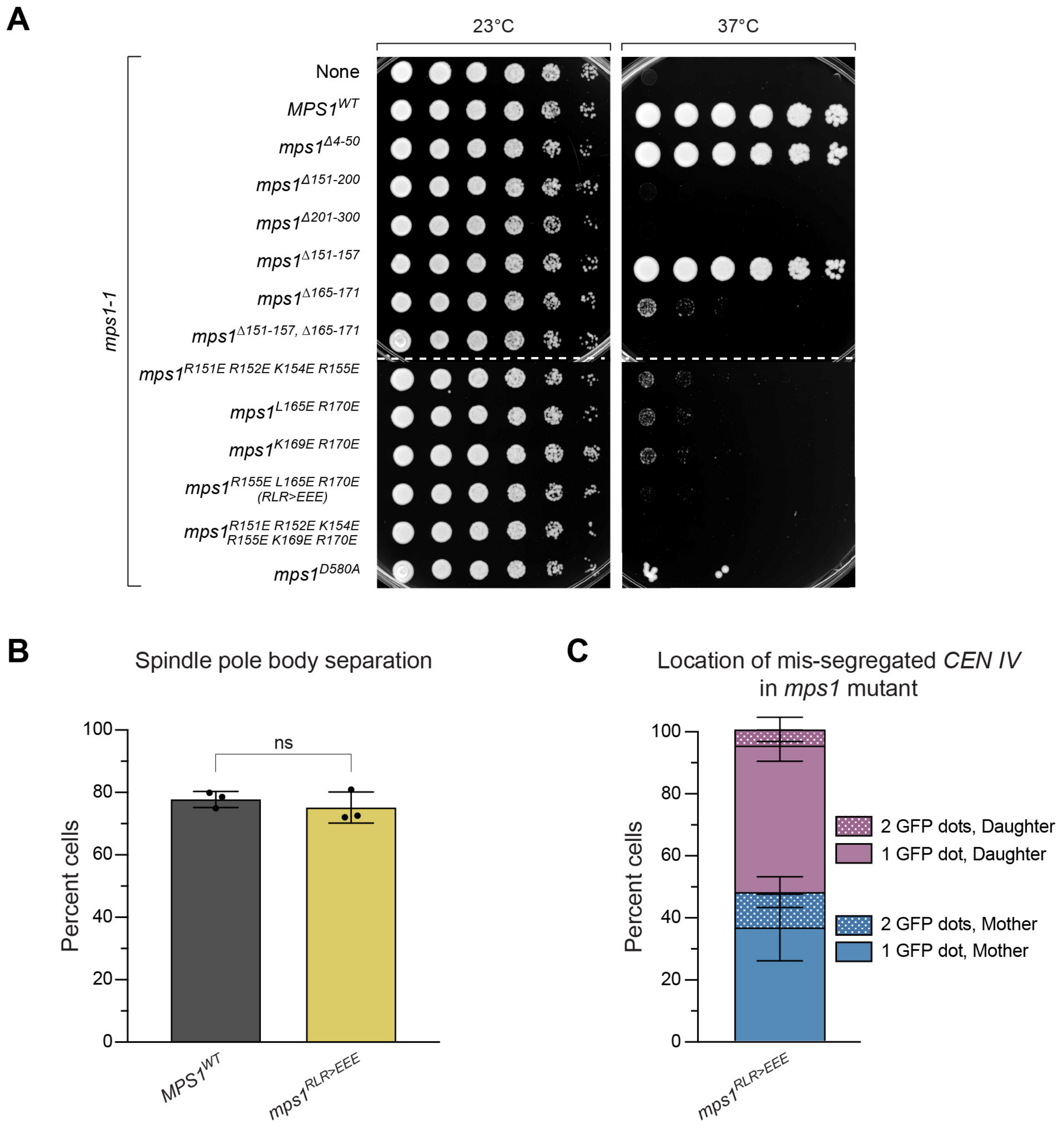
A
23°C
37°C
mps1-1
None
MPS1WT
mps1Δ4-50
mps1Δ151-200
mps1Δ201-300
mps1Δ151-157
mps1Δ165-171
mps1Δ151-157, Δ165-171
mps1R151E R152E K154E R155E
mps1L165E R170E
mps1K169E R170E
mps1R155E L165E R170E (RLR>EEE)
mps1R151E R152E K154E R155E K169E R170E
mps1D580A
B
Spindle pole body separation
Percent cells
100
80
60
40
20
0
ns
MPS1WT
mps1RLR>EEE
C
Location of mis-segregated CEN IV in mps1 mutant
Percent cells
100
80
60
40
20
0
2 GFP dots, Daughter
1 GFP dot, Daughter
2 GFP dots, Mother
1 GFP dot, Mother
mps1RLR>EEE

**Figure S4. The *mps1*$^{RLR>EEE}$ mutant displays normal spindle pole body separation but aberrant localization of *CEN IV* GFP.**

(A) Complementation of the temperature sensitive *mps1-1* allele (M56) by ectopically expressed *MPS1* (*MPS1*$^{WT}$, M4407; *mps1*$^{\Delta 4\text{-}50}$, M4417; *mps1*$^{\Delta 151\text{-}200}$, M4412; *mps1*$^{\Delta 201\text{-}300}$, M4413; *mps1*$^{\Delta 151\text{-}157}$, M4409; *mps1*$^{\Delta 165\text{-}171}$, M4410; *mps1*$^{\Delta 151\text{-}157,\ 165\text{-}171}$, M4418; *mps1*$^{R151E\ R152E\ K154E\ R155E}$, M4415; *mps1*$^{L165E\ R170E}$, M4414; *mps1*$^{K169E\ R170E}$, M4416; *mps1*$^{R155E\ L165E\ R170E\ (RLR>EEE)}$, M4949; *mps1*$^{R151E\ R152E\ K154E\ R155E\ K169E\ R170E}$, M4420; *mps1*$^{D580A}$ kinase dead, M4411). Cells were serially diluted five-fold, spotted onto plates, and grown at *mps1-1* permissive (23°C) and non-permissive (37°C) temperatures. We note that in our strain background, *mps1*$^{\Delta 4\text{-}50}$ supports viability, which contrasts with previous findings[57].

(B) Quantitation of percent of cells with spindle pole body separation in *MPS1*$^{WT}$ (M4714) and *mps1*$^{RLR>EEE}$ (M4710, M4947) cells from Figure 4D. Strains also carry the *mps1-1* temperature sensitive allele, *CEN IV* marked with GFP (*CEN IV:lacO LacI-GFP*), and *SPC110-mCherry* (spindle pole marker). Exponentially growing cells were arrested with 1 µg/mL alpha factor for 3 hours, followed by release into fresh media at 37°C for 1.5 hours. Error bars indicate the standard deviation among three replicates (n = 105-139 cells for each replicate). Significance was determined by a two-tailed unpaired *t* test (ns; *P* = 0.465).

**(C)** Quantitation of the location of mis-localized *CEN IV* GFP from *MPS1*$^{WT}$ (M4714) and *mps1*$^{RLR>EEE}$ (M4710, M4947, M4948), shown in Figure 4D. Due to the difference in phenotypes between *nuf2* mutants and *mps1*$^{RLR>EEE}$, three independent strains were analyzed to ensure validity of the result. The percent of cells with 1 vs. 2 *CEN IV* GFP signals localized to the mother or the daughter is shown. Error bars indicate the standard deviation among 4 replicates.

### Figure S5

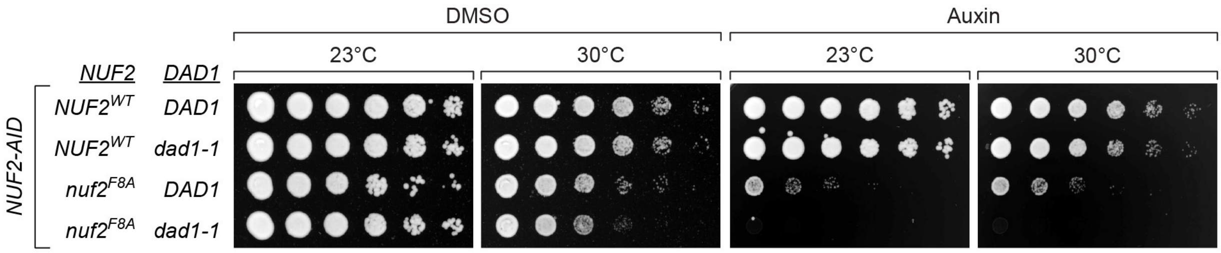

DMSO
Auxin
23°C
30°C
23°C
30°C
NUF2
DAD1
NUF2-AID
NUF2WT DAD1
NUF2WT dad1-1
nuf2F8A DAD1
nuf2F8A dad1-1

**Figure S5. A temperature sensitive *dad1-1* allele is synthetic lethal with *nuf2* mutants.** Cell viability assay assessing synthetic lethality of a *nuf2* mutant allele with the temperature sensitive *dad1-1* allele (extension of Fig 5C). Strains carry a *NUF2-AID* allele and ectopic copies of *NUF2-3HA* (*$NUF2^{WT}$ DAD1*, M2038; *$NUF2^{WT}$ dad1-1*, M4465; *$nuf2^{F8A}$ DAD1*, M2151; *$nuf2^{F8A}$ dad1-1*, M4924). Cells were serially diluted, spotted onto plates, and grown at *dad-1* permissive (23°C) and semi-permissive (30°C) temperatures.

Figure S6

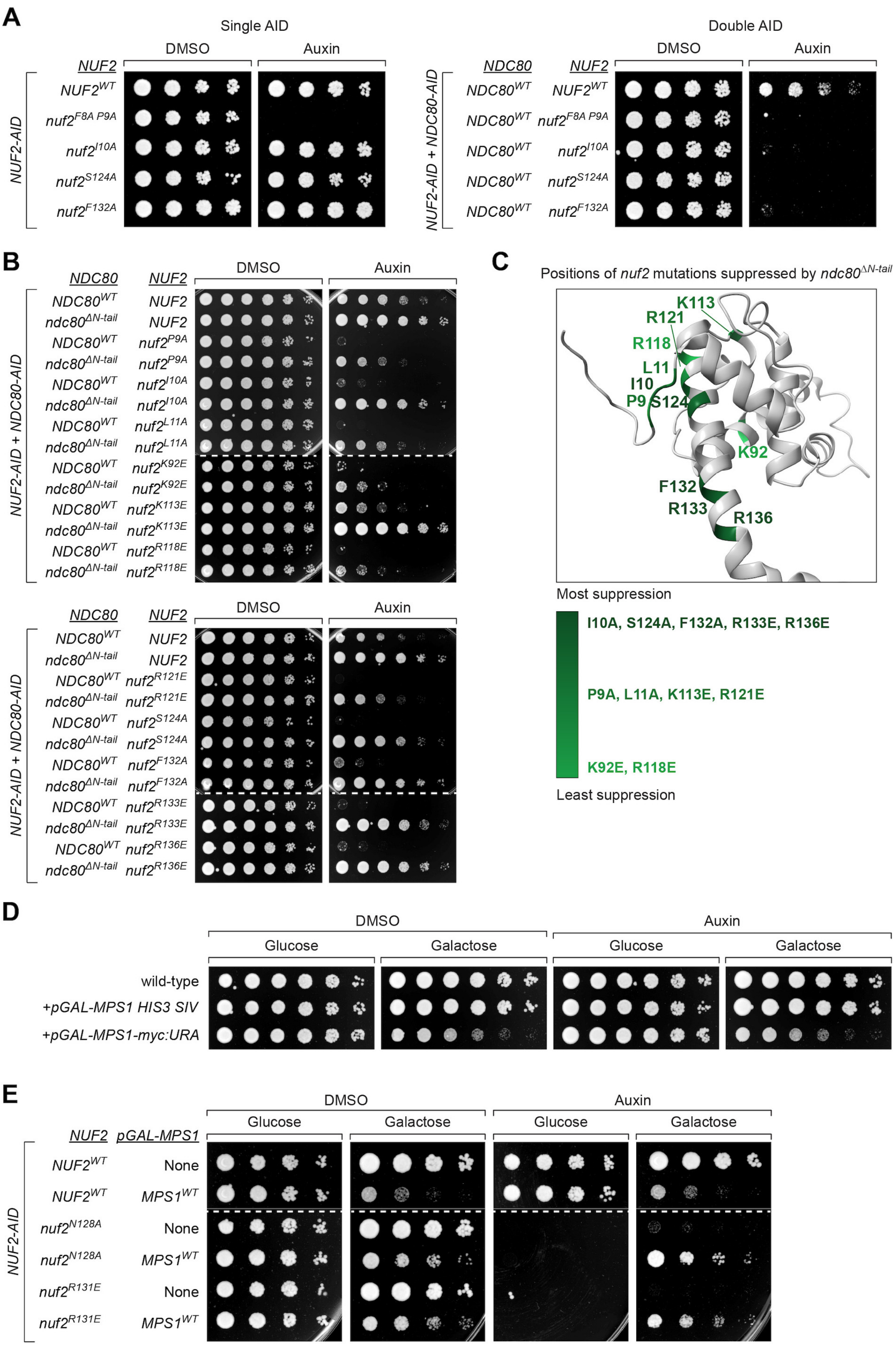

A
Single AID
DMSO
Auxin
NUF2
NUF2-AID
NUF2WT
nuf2F8A P9A
nuf2I10A
nuf2S124A
nuf2F132A
Double AID
NDC80
NUF2-AID + NDC80-AID
NDC80WT
B
ndc80ΔN-tail
nuf2P9A
nuf2L11A
nuf2K92E
nuf2K113E
nuf2R118E
nuf2R121E
nuf2R133E
nuf2R136E
C
Positions of nuf2 mutations suppressed by ndc80ΔN-tail
K113
R121
R118
L11
I10
P9
S124
K92
F132
R133
R136
Most suppression
I10A, S124A, F132A, R133E, R136E
P9A, L11A, K113E, R121E
K92E, R118E
Least suppression
D
Glucose
Galactose
wild-type
+pGAL-MPS1 HIS3 SIV
+pGAL-MPS1-myc:URA
E
pGAL-MPS1
None
MPS1WT
nuf2N128A
nuf2R131E

### Figure S6

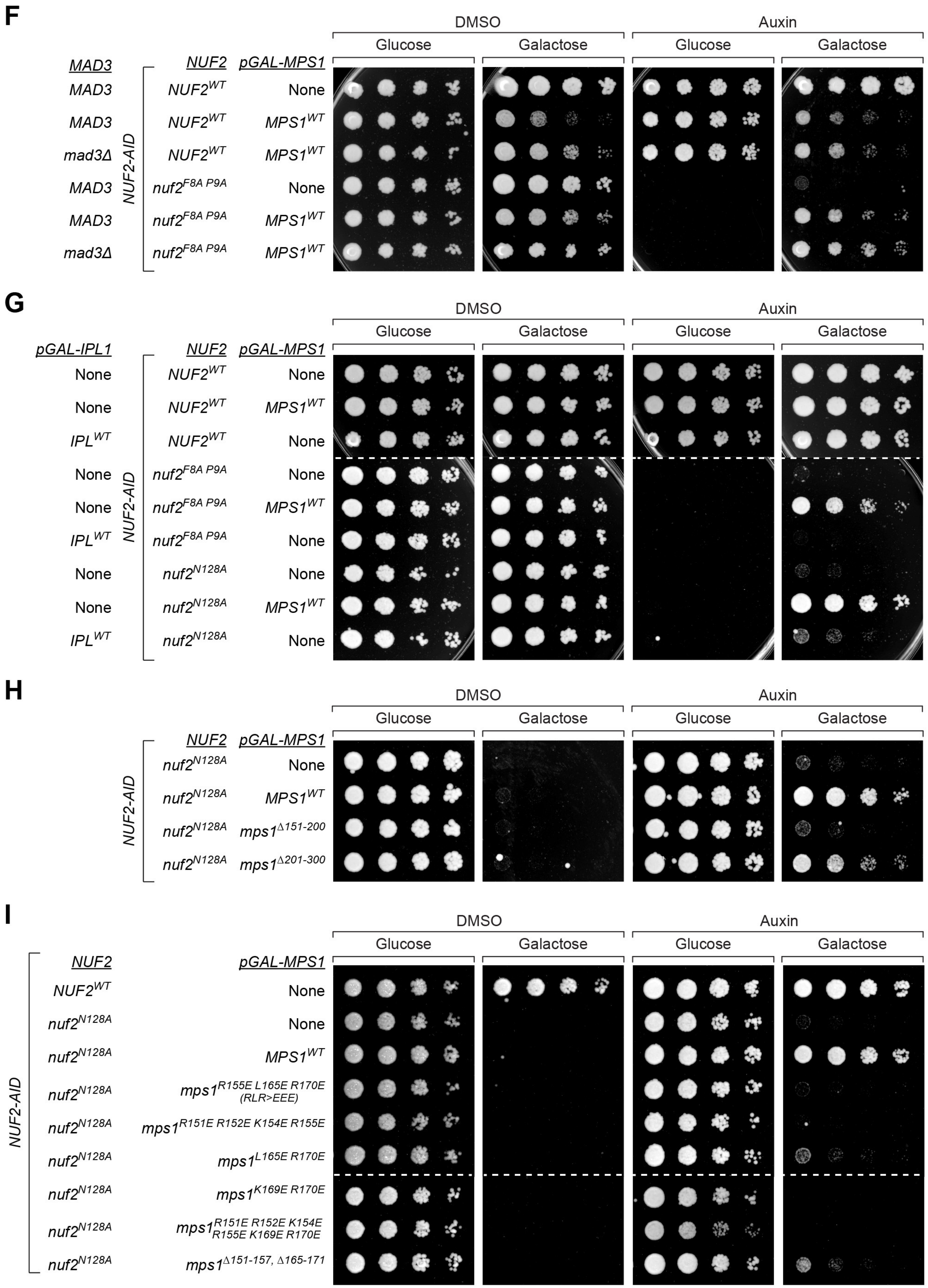

F
DMSO
Auxin
Glucose
Galactose
Glucose
Galactose
MAD3
NUF2
pGAL-MPS1
NUF2-AID
MAD3 $NUF2^{WT}$ None
MAD3 $NUF2^{WT}$ $MPS1^{WT}$
mad3Δ $NUF2^{WT}$ $MPS1^{WT}$
MAD3 $nuf2^{F8A\ P9A}$ None
MAD3 $nuf2^{F8A\ P9A}$ $MPS1^{WT}$
mad3Δ $nuf2^{F8A\ P9A}$ $MPS1^{WT}$
G
DMSO
Auxin
Glucose
Galactose
Glucose
Galactose
pGAL-IPL1
NUF2
pGAL-MPS1
NUF2-AID
None $NUF2^{WT}$ None
None $NUF2^{WT}$ $MPS1^{WT}$
$IPL^{WT}$ $NUF2^{WT}$ None
None $nuf2^{F8A\ P9A}$ None
None $nuf2^{F8A\ P9A}$ $MPS1^{WT}$
$IPL^{WT}$ $nuf2^{F8A\ P9A}$ None
None $nuf2^{N128A}$ None
None $nuf2^{N128A}$ $MPS1^{WT}$
$IPL^{WT}$ $nuf2^{N128A}$ None
H
DMSO
Auxin
Glucose
Galactose
Glucose
Galactose
NUF2
pGAL-MPS1
NUF2-AID
$nuf2^{N128A}$ None
$nuf2^{N128A}$ $MPS1^{WT}$
$nuf2^{N128A}$ $mps1^{\Delta 151-200}$
$nuf2^{N128A}$ $mps1^{\Delta 201-300}$
I
DMSO
Auxin
Glucose
Galactose
Glucose
Galactose
NUF2
pGAL-MPS1
NUF2-AID
$NUF2^{WT}$ None
$nuf2^{N128A}$ None
$nuf2^{N128A}$ $MPS1^{WT}$
$nuf2^{N128A}$ $mps1^{R155E\ L165E\ R170E}$ (RLR>EEE)
$nuf2^{N128A}$ $mps1^{R151E\ R152E\ K154E\ R155E}$
$nuf2^{N128A}$ $mps1^{L165E\ R170E}$
$nuf2^{N128A}$ $mps1^{K169E\ R170E}$
$nuf2^{N128A}$ $mps1^{R151E\ R152E\ K154E\ R155E\ K169E\ R170E}$
$nuf2^{N128A}$ $mps1^{\Delta 151-157,\ \Delta 165-171}$

**Figure S6. Suppression of *nuf2* mutants by deletion of the Ndc80 N-terminal tail and over-expression of Mps1.**

(A) Yeast cell viability assay, comparing ectopic *nuf2* mutants expressed in a "single AID" strain with *NUF2-AID* (*NUF2*$^{WT}$, M2038; *nuf2*$^{F8A\ P9A}$, M2042; *nuf2*$^{I10A}$, M2191; *nuf2*$^{S124A}$, M2040; *nuf2*$^{F132A}$, M2545) to those in a "double AID" strain containing both *NUF2-AID* and *NDC80-AID* alleles, as well as an ectopic copy of *NDC80*$^{WT}$ (*NUF2*$^{WT}$, M2998; *nuf2*$^{F8A\ P9A}$, M2643; *nuf2*$^{I10A}$, M2654; *nuf2*$^{S124A}$, M2641; *nuf2*$^{F132A}$, M2841). Cells were serially diluted five-fold and spotted onto plates containing DMSO or 250 μM auxin. Note that the "double AID" strain is genetically sensitized such that mutants with no detectable phenotype in the "single AID" strain show large growth defects in the "double AID" genetic background.

(B) Yeast cell viability assay comparing *nuf2* mutant phenotypes in *NDC80*$^{WT}$ and *ndc80*$^{\Delta N\text{-}tail}$ strains. Double *NUF2-AID*, *NDC80-AID* strains carry ectopic copies of *NUF2-3HA* and *NDC80*$^{WT}$*-3HA* or *NDC80*$^{\Delta N\text{-}tail}$*-3HA*, respectively (*NUF2*$^{WT}$, M2998, M2999; *nuf2*$^{P9A}$, M2647, M2748; *nuf2*$^{I10A}$, M2654, M2755; *nuf2*$^{L11A}$, M2655, M2756; *nuf2*$^{K92E}$, M2666, M2763; *nuf2*$^{K113E}$, M2648, M2749; *nuf2*$^{R118E}$, M2649, M2750; *nuf2*$^{R121E}$, M2660, M2757; *nuf2*$^{S124A}$, M2641, M2742; *nuf2*$^{F132A}$, M2841, M2774; *nuf2*$^{R133E}$, M2662, M2759; *nuf2*$^{R136E}$, M2663, M2760). Cells were serially diluted five-fold and spotted onto plates containing DMSO or 250 μM auxin.

(C) Structure of *S. cerevisiae* Nuf2 (5TCS) illustrating the residues with mutants that are suppressed when combined with an *ndc80*$^{N\text{-}tail}$ allele in the "double AID" strain containing *NUF2-AID* and *NDC80-AID*.

(D) Yeast cell viability assay to examine the effect of *MPS1* overexpression on cell viability. Wild-type (M3), *pGAL-MPS1* as a single integration vector at the *HIS3* locus (M4618) and *pGAL-MPS1-myc* from a pRS vector (potentially multiple copies) integrated at the *URA3* locus (M2850) were serially diluted five-fold and spotted onto plates containing either glucose or galactose. Note that these observations suggest the levels of *MPS1* overexpression are not sufficient to lead to a constitutive SAC induced arrest, which is not viable[65].

(E) Yeast cell viability assay to assess the effect of *pGAL*-driven overexpression of Mps1 in an additional *nuf2* mutant. Yeast cell viability assay in strains containing *NUF2-AID* and ectopic copies of *NUF2-3HA,* without or with an ectopic *pGAL-MPS1-myc* allele at the *URA3* locus (*NUF2*$^{WT}$, M2038, M2680; *nuf2*$^{N128A}$, M2414, M2709; *nuf2*$^{R131E}$, M2201,

M2697). Cells were serially diluted five-fold and spotted onto plates containing either glucose or galactose, as well as DMSO or 250 μM auxin.

(F) Yeast cell viability assay to assess *pGAL-MPS1* suppression of *nuf2* mutants in a SAC mutant. Strains containing *NUF2-AID* and ectopic copies of *NUF2-3HA,* without or with an ectopic *pGAL-MPS1-myc* allele at the *URA3* locus in both *MAD3* and *mad3*Δ backgrounds are shown (*NUF2*$^{WT}$, with no *pGAL-MPS1* M2038, with *pGAL-MPS1*$^{WT}$ M2680, with *pGAL-MPS1*$^{WT}$ *mad3*Δ M2867; *nuf2*$^{F8A\ P9A}$, with no *pGAL-MPS1* M2042, with *pGAL-MPS1*$^{WT}$ 2683, with *pGAL-MPS1*$^{WT}$ *mad3*Δ M2870). Cells were serially diluted and spotted onto plates containing either glucose or galactose, as well as DMSO or 250 μM auxin.

(G) Yeast cell viability assay to compare growth of *nuf2* mutant strains overexpressing either ectopic *pGAL-MPS1* or *pGAL-IPL1*. Strains contain *NUF2-AID* and ectopic copies of *NUF2-3HA* with the addition of either *pGAL-MPS1* or *pGAL-IPL1* as single copies integrated at the *HIS3* locus (*NUF2*$^{WT}$, M2038; *NUF2*$^{WT}$ with *pGAL-MPS1*, M3586; *NUF2*$^{WT}$ with *pGAL-IPL1*, M4171; *nuf2*$^{F8A\ P9A}$, M2042; *nuf2*$^{F8A\ P9A}$ with *pGAL-MPS1*, M3587; *nuf2*$^{F8A\ P9A}$ with *pGAL-IPL1*, M4172; *nuf2*$^{N128A}$, M2414; *nuf2*$^{N128A}$ with *pGAL-MPS1*, M3588; *nuf2*$^{N128A}$ with *pGAL-IPL1*, M4173). Cells were serially diluted and spotted onto plates containing either glucose or galactose, as well as DMSO or 250 μM auxin.

(H) Yeast cell viability assay to compare the effect of overexpression of *pGAL-MPS1*$^{WT}$ with *pGAL-mps1* deletions on the growth phenotype of *nuf2*$^{N128A}$. Strains carry *NUF2-AID*, an ectopic copy of *NUF2*$^{WT}$*-3HA* or *nuf2*$^{N128A}$*-3HA,* and a single, ectopic *pGAL-MPS1* allele (*NUF2*$^{WT}$, M2038; *nuf2*$^{N128A}$ with no *pGAL-MPS1* allele, "None", M2414; with *pGAL-MPS1*$^{WT}$, M3587; with *pGAL-mps1*$^{\Delta 151-200}$, M4055; with *pGAL-mps1*$^{\Delta 201-300}$, M4056). A previous study implicated Mps1 residues 151-200 in biorientation and residues 201-300 in spindle pole body duplication[57].

(I) Yeast cell viability assay to compare the level of *nuf2*$^{N128A}$ suppression by *pGAL-mps1*$^{RLR>EEE}$ overexpression with that of other *mps1* mutants at the Nuf2-Mps1 interface. Strains contain *NUF2-AID*, an ectopic copy of *NUF2*$^{WT}$ *or nuf2*$^{N128A}$*,* and a single, ectopic *pGAL-MPS1* allele (*NUF2*$^{WT}$, M2038; *nuf2*$^{N128A}$ with no *pGAL-MPS1* allele, M2414; with *pGAL-MPS1*$^{WT}$, M3587; *nuf2*$^{N128A}$ with *mps1*$^{RLR>EEE}$, M4946; *nuf2*$^{N128A}$ with *mps1*$^{R151E\ R152E\ K154E\ R155E}$, M4641; *nuf2*$^{N128A}$ with *mps1*$^{L165E\ R170E}$, M4640; *nuf2*$^{N128A}$ with *mps1*$^{K169E\ R170E}$, M4642; *nuf2*$^{N128A}$ with *mps1*$^{R151E\ R152E\ K154E\ R155E\ K169E\ R170E}$, M4645; *nuf2*$^{N128A}$ with *mps1*$^{\Delta151\text{-}157,\Delta165\text{-}171}$, M4643). Cells were serially diluted five-fold and spotted onto plates containing either glucose or galactose, as well as DMSO or 250 µM auxin.

**Table S1. All strains used in this study are derivatives of M3 (W303).**

| Strain | Relevant Genotype | Figure |
|---|---|---|
| M3 (W303) | *MAT***a** *ura3-1 leu2-3,112 his3-11 trp1-1 can1-100 ade2-1 bar1-1* | S6D |
| M35 | *MAT***a** *mad2Δ::URA3* | 3A |
| M56 | *MAT***a** *mps1-1* | 4C, S4A |
| M457 | *MAT***a** *DSN1-HIS-FLAG:URA3 NDC80-3HA:KanMX DAM1-myc9:TRP1* | 6F |
| M469 | *MAT***a** *DSN1-HIS-FLAG:URA3 NDC80-3HA:KanMX DAM1-myc9:TRP1 mps1-1* | 6F |
| M692 | *MAT***a** *NDC80-3V5-IAA7:KanMX leu2::pGPD1-OsTIR1:LEU2 trp1::pNDC80-NDC80-3HA:TRP1* | 2A, S2 |
| M693 | *MAT***a** *NDC80-3V5-IAA7:KanMX leu2::pGPD1-OsTIR1:LEU2 trp1::pNDC80-ndc80(K204E)-3HA:TRP1* | 2A, S2 |
| M694 | *MAT***a** *NDC80-3V5-IAA7:KanMX leu2::pGPD1-OsTIR1:LEU2 trp1::pNDC80-ndc80(K122E K204E)-3HA:TRP1* | 2A,B |
| M1375 | *MAT***a** *TOR1-1 fpr1Δ::NatMX* | 7B |
| M1463 | *MAT***a** *TOR1-1 fpr1Δ::NatMX NDC80-FKBP12:His3MX MPS1-FRB:KanMX DSN1-HIS-FLAG:URA3* | 7B |
| M1889 | *MAT***a** *NUF2-3V5-IAA7:KanMX trp1::pGPD1-OsTIR1:TRP1 DSN1-HIS-FLAG:URA3* | 1B, 2A, S1C,D |
| M2038 | *MAT***a** *NUF2-3V5-IAA7:KanMX trp1::pGPD1-OsTIR1:TRP1 DSN1-HIS-FLAG:URA3 leu2::pNUF2-NUF2-3HA:LEU2* | 1B, 2A, 3A, 5A, 6C,E, S1C,D, S2, S5, S6A,E, F,G,H,I |
| M2040 | *MAT***a** *NUF2-3V5-IAA7:KanMX trp1::pGPD1-OsTIR1:TRP1 DSN1-HIS-FLAG:URA3 leu2::pNUF2-nuf2(S124A)-3HA:LEU2* | 1B, S1C,D, S6A |
| M2041 | *MAT***a** *NUF2-3V5-IAA7:KanMX trp1::pGPD1-OsTIR1:TRP1 DSN1-HIS-FLAG:URA3 leu2::pNUF2-nuf2(S124D)-3HA:LEU2* | 1B, 6C, S1C,D |
| M2042 | *MAT***a** *NUF2-3V5-IAA7:KanMX trp1::pGPD1-OsTIR1:TRP1 DSN1-HIS-FLAG:URA3 leu2::pNUF2-nuf2(F8A P9A)-3HA:LEU2* | 1B, 2A, 3A, 6C, S1C,D, S6A,F, G |
| M2043 | *MAT***a** *NUF2-3V5-IAA7:KanMX trp1::pGPD1-OsTIR1:TRP1 DSN1-HIS-FLAG:URA3 leu2::pNUF2-nuf2(S2A R3A N4A Q5A)-3HA:LEU2* | S1D |
| M2044 | *MAT***a** *NUF2-3V5-IAA7:KanMX trp1::pGPD1-OsTIR1:TRP1 DSN1-HIS-FLAG:URA3 leu2::pNUF2-nuf2(D6A V7A I10A L11A)-3HA:LEU2* | S1D |
| M2045 | *MAT***a** *NUF2-3V5-IAA7:KanMX trp1::pGPD1-OsTIR1:TRP1 DSN1-HIS-FLAG:URA3 leu2::pNUF2-nuf2(Δ2-11)-3HA:LEU2* | S1D |
| M2046 | *MAT***a** *NUF2-3V5-IAA7:KanMX trp1::pGPD1-OsTIR1:TRP1 DSN1-HIS-FLAG:URA3 leu2::pNUF2-nuf2(Δ6-7)-3HA:LEU2* | S1D |
| M2047 | *MAT***a** *NUF2-3V5-IAA7:KanMX trp1::pGPD1-OsTIR1:TRP1 DSN1-HIS-FLAG:URA3 leu2::pNUF2-nuf2(Δ8-11)-3HA:LEU2* | S1D |

| | | |
|---|---|---|
| M2048 | *MAT**a** NUF2-3V5-IAA7:KanMX trp1::pGPD1-OsTIR1:TRP1 DSN1-HIS-FLAG:URA3 leu2::pNUF2-nuf2(Δ2-5)-3HA:LEU2* | S1D |
| M2151 | *MAT**a** NUF2-3V5-IAA7:KanMX trp1::pGPD1-OsTIR1:TRP1 DSN1-HIS-FLAG:URA3 leu2::pNUF2-nuf2(F8A)-3HA:LEU2* | S1D, S5 |
| M2152 | *MAT**a** NUF2-3V5-IAA7:KanMX trp1::pGPD1-OsTIR1:TRP1 DSN1-HIS-FLAG:URA3 leu2::pNUF2-nuf2(P9A)-3HA:LEU2* | 5B, S1D |
| M2153 | *MAT**a** NUF2-3V5-IAA7:KanMX trp1::pGPD1-OsTIR1:TRP1 DSN1-HIS-FLAG:URA3 leu2::pNUF2-nuf2(K113E)-3HA:LEU2* | S1D |
| M2154 | *MAT**a** NUF2-3V5-IAA7:KanMX trp1::pGPD1-OsTIR1:TRP1 DSN1-HIS-FLAG:URA3 leu2::pNUF2-nuf2(R118E)-3HA:LEU2* | S1D |
| M2191 | *MAT**a** NUF2-3V5-IAA7:KanMX trp1::pGPD1-OsTIR1:TRP1 DSN1-HIS-FLAG:URA3 leu2::pNUF2-nuf2(I10A)-3HA:LEU2* | S1D, S6A |
| M2192 | *MAT**a** NUF2-3V5-IAA7:KanMX trp1::pGPD1-OsTIR1:TRP1 DSN1-HIS-FLAG:URA3 leu2::pNUF2-nuf2(L11A)-3HA:LEU2* | S1D |
| M2200 | *MAT**a** NUF2-3V5-IAA7:KanMX trp1::pGPD1-OsTIR1:TRP1 DSN1-HIS-FLAG:URA3 leu2::pNUF2-nuf2(R121E)-3HA:LEU2* | S1D |
| M2201 | *MAT**a** NUF2-3V5-IAA7:KanMX trp1::pGPD1-OsTIR1:TRP1 DSN1-HIS-FLAG:URA3 leu2::pNUF2-nuf2(R131E)-3HA:LEU2* | S1D, S6E |
| M2202 | *MAT**a** NUF2-3V5-IAA7:KanMX trp1::pGPD1-OsTIR1:TRP1 DSN1-HIS-FLAG:URA3 leu2::pNUF2-nuf2(R133E)-3HA:LEU2* | S1D |
| M2203 | *MAT**a** NUF2-3V5-IAA7:KanMX trp1::pGPD1-OsTIR1:TRP1 DSN1-HIS-FLAG:URA3 leu2::pNUF2-nuf2(R136E)-3HA:LEU2* | S1D |
| M2227 | *MAT**a** NUF2-3V5-IAA7:KanMX trp1::pGPD1-OsTIR1:TRP1 DSN1-HIS-FLAG:URA3 leu2::pNUF2-nuf2(R35E)-3HA:LEU2* | S1D |
| M2228 | *MAT**a** NUF2-3V5-IAA7:KanMX trp1::pGPD1-OsTIR1:TRP1 DSN1-HIS-FLAG:URA3 leu2::pNUF2-nuf2(K46E)-3HA:LEU2* | S1D |
| M2229 | *MAT**a** NUF2-3V5-IAA7:KanMX trp1::pGPD1-OsTIR1:TRP1 DSN1-HIS-FLAG:URA3 leu2::pNUF2-nuf2(K92E)-3HA:LEU2* | S1D |
| M2230 | *MAT**a** NUF2-3V5-IAA7:KanMX trp1::pGPD1-OsTIR1:TRP1 DSN1-HIS-FLAG:URA3 leu2::pNUF2-nuf2(K96E)-3HA:LEU2* | S1D |
| M2262 | *MAT**a** NUF2-3V5-IAA7:KanMX trp1::pGPD1-OsTIR1:TRP1 DSN1-HIS-FLAG:URA3 leu2::pNUF2-nuf2(F8A P9A L11A)-3HA:LEU2* | S1D |
| M2263 | *MAT**a** NUF2-3V5-IAA7:KanMX trp1::pGPD1-OsTIR1:TRP1 DSN1-HIS-FLAG:URA3 leu2::pNUF2-nuf2(R118E R133E)-3HA:LEU2* | S1D |
| M2264 | *MAT**a** NUF2-3V5-IAA7:KanMX trp1::pGPD1-OsTIR1:TRP1 DSN1-HIS-FLAG:URA3 leu2::pNUF2-nuf2(R118E R121E)-3HA:LEU2* | S1D |
| M2323 | *MAT**a** NUF2-3V5-IAA7:KanMX trp1::pGPD1-OsTIR1:TRP1 DSN1-HIS-FLAG:URA3 leu2::pNUF2-nuf2(K113E R118E)-3HA:LEU2* | S1D |
| M2413 | *MAT**a** NUF2-3V5-IAA7:KanMX trp1::pGPD1-OsTIR1:TRP1 DSN1-HIS-FLAG:URA3 leu2::pNUF2-nuf2(D105K)-3HA:LEU2* | S1D |
| M2414 | *MAT**a** NUF2-3V5-IAA7:KanMX trp1::pGPD1-OsTIR1:TRP1 DSN1-HIS-FLAG:URA3 leu2::pNUF2-nuf2(N128A)-3HA:LEU2* | 1B, 2A, 3A, 6C,E, S1C,D, S6E,G H,I |
| M2544 | *MAT**a** NUF2-3V5-IAA7:KanMX trp1::pGPD1-OsTIR1:TRP1 DSN1-HIS-FLAG:URA3 leu2::pNUF2-nuf2(D110K)-3HA:LEU2* | S1D |
| M2545 | *MAT**a** NUF2-3V5-IAA7:KanMX trp1::pGPD1-OsTIR1:TRP1 DSN1-HIS-FLAG:URA3 leu2::pNUF2-nuf2(F132A)-3HA:LEU2* | S1D, S6A |

| | | |
|---|---|---|
| M2630 | *MAT***a** *DSN1-HIS-FLAG:URA3* | 4A |
| M2633 | *MAT***a** *NUF2-3V5-IAA7:KanMX trp1::pGPD1-OsTIR1:TRP1 DSN1-HIS-FLAG:URA3 leu2::pNUF2-nuf2(A125D)-3HA:LEU2* | S1D |
| M2641 | *MATα NUF2-3V5-IAA7:KanMX NDC80-3V5-IAA7:KanMX his3::pGPD1-OsTIR1:HIS3 leu2::pNUF2-nuf2(S124A)-3HA:LEU2 trp1::pNDC80-NDC80-3HA:TRP1* | 6B, S6A,B |
| M2643 | *MATα NUF2-3V5-IAA7:KanMX NDC80-3V5-IAA7:KanMX his3::pGPD1-OsTIR1:HIS3 leu2::pNUF2-nuf2(F8A P9A)-3HA:LEU2 trp1::pNDC80-NDC80-3HA:TRP1* | S6A |
| M2647 | *MATα NUF2-3V5-IAA7:KanMX NDC80-3V5-IAA7:KanMX his3::pGPD1-OsTIR1:HIS3 leu2::pNUF2-nuf2(P9A)-3HA:LEU2 trp1::pNDC80-NDC80-3HA:TRP1* | S6B |
| M2648 | *MATα NUF2-3V5-IAA7:KanMX NDC80-3V5-IAA7:KanMX his3::pGPD1-OsTIR1:HIS3 leu2::pNUF2-nuf2(K113E)-3HA:LEU2 trp1::pNDC80-NDC80-3HA:TRP1* | S6B |
| M2649 | *MATα NUF2-3V5-IAA7:KanMX NDC80-3V5-IAA7:KanMX his3::pGPD1-OsTIR1:HIS3 leu2::pNUF2-nuf2(R118E)-3HA:LEU2 trp1::pNDC80-NDC80-3HA:TRP1* | S6B |
| M2654 | *MATα NUF2-3V5-IAA7:KanMX NDC80-3V5-IAA7:KanMX his3::pGPD1-OsTIR1:HIS3 leu2::pNUF2-nuf2(I10A)-3HA:LEU2 trp1::pNDC80-NDC80-3HA:TRP1* | 6B, S6A,B |
| M2655 | *MATα NUF2-3V5-IAA7:KanMX NDC80-3V5-IAA7:KanMX his3::pGPD1-OsTIR1:HIS3 leu2::pNUF2-nuf2(L11A)-3HA:LEU2 trp1::pNDC80-NDC80-3HA:TRP1* | S6B |
| M2660 | *MATα NUF2-3V5-IAA7:KanMX NDC80-3V5-IAA7:KanMX his3::pGPD1-OsTIR1:HIS3 leu2::pNUF2-nuf2(R121E)-3HA:LEU2 trp1::pNDC80-NDC80-3HA:TRP1* | S6B |
| M2662 | *MATα NUF2-3V5-IAA7:KanMX NDC80-3V5-IAA7:KanMX his3::pGPD1-OsTIR1:HIS3 leu2::pNUF2-nuf2(R133E)-3HA:LEU2 trp1::pNDC80-NDC80-3HA:TRP1* | S6B |
| M2663 | *MATα NUF2-3V5-IAA7:KanMX NDC80-3V5-IAA7:KanMX his3::pGPD1-OsTIR1:HIS3 leu2::pNUF2-nuf2(R136E)-3HA:LEU2 trp1::pNDC80-NDC80-3HA:TRP1* | S6B |
| M2666 | *MATα NUF2-3V5-IAA7:KanMX NDC80-3V5-IAA7:KanMX his3::pGPD1-OsTIR1:HIS3 leu2::pNUF2-nuf2(K92E)-3HA:LEU2 trp1::pNDC80-NDC80-3HA:TRP1* | S6B |
| M2680 | *MAT***a** *NUF2-3V5-IAA7:KanMX his3::pGPD1-OsTIR1:HIS3 DSN1-HIS-FLAG:URA3 leu2::pNUF2-NUF2-3HA:LEU2 ura3::pGAL-MPS1-myc:URA3* | S6E,F |
| M2683 | *MAT***a** *NUF2-3V5-IAA7:KanMX his3::pGPD1-OsTIR1:HIS3 DSN1-HIS-FLAG:URA3 leu2::pNUF2-nuf2(F8A P9A)-3HA:LEU2 ura3::pGAL-MPS1-myc:URA3* | S6F |
| M2697 | *MAT***a** *NUF2-3V5-IAA7:KanMX his3::pGPD1-OsTIR1:HIS3 DSN1-HIS-FLAG:URA3 leu2::pNUF2-nuf2(R131E)-3HA:LEU2 ura3::pGAL-MPS1-myc:URA3* | S6E |
| M2709 | *MAT***a** *NUF2-3V5-IAA7:KanMX his3::pGPD1-OsTIR1:HIS3 DSN1-HIS-FLAG:URA3 leu2::pNUF2-nuf2(N128A)-3HA:LEU2 ura3::pGAL-MPS1-myc:URA3* | S6E |

| | | |
|---|---|---|
| M2742 | *MATα NUF2-3V5-IAA7:KanMX NDC80-3V5-IAA7:KanMX his3::pGPD1-OsTIR1:HIS3 leu2::pNUF2-nuf2(S124A)-3HA:LEU2 trp1::pNDC80-ndc80(ΔN-tail)-3HA:TRP1* | 6B, S6B |
| M2748 | *MATα NUF2-3V5-IAA7:KanMX NDC80-3V5-IAA7:KanMX his3::pGPD1-OsTIR1:HIS3 leu2::pNUF2-nuf2(P9A)-3HA:LEU2 trp1::pNDC80-ndc80(ΔN-tail)-3HA:TRP1* | S6B |
| M2749 | *MATα NUF2-3V5-IAA7:KanMX NDC80-3V5-IAA7:KanMX his3::pGPD1-OsTIR1:HIS3 leu2::pNUF2-nuf2(K113E)-3HA:LEU2 trp1::pNDC80-ndc80(ΔN-tail)-3HA:TRP1* | S6B |
| M2750 | *MATα NUF2-3V5-IAA7:KanMX NDC80-3V5-IAA7:KanMX his3::pGPD1-OsTIR1:HIS3 leu2::pNUF2-nuf2(R118E)-3HA:LEU2 trp1::pNDC80-ndc80(ΔN-tail)-3HA:TRP1* | S6B |
| M2755 | *MATα NUF2-3V5-IAA7:KanMX NDC80-3V5-IAA7:KanMX his3::pGPD1-OsTIR1:HIS3 leu2::pNUF2-nuf2(I10A)-3HA:LEU2 trp1::pNDC80-ndc80(ΔN-tail)-3HA:TRP1* | 6B, S6B |
| M2756 | *MATα NUF2-3V5-IAA7:KanMX NDC80-3V5-IAA7:KanMX his3::pGPD1-OsTIR1:HIS3 leu2::pNUF2-nuf2(L11A)-3HA:LEU2 trp1::pNDC80-ndc80(ΔN-tail)-3HA:TRP1* | S6B |
| M2757 | *MATα NUF2-3V5-IAA7:KanMX NDC80-3V5-IAA7:KanMX his3::pGPD1-OsTIR1:HIS3 leu2::pNUF2-nuf2(R121E)-3HA:LEU2 trp1::pNDC80-ndc80(ΔN-tail)-3HA:TRP1* | S6B |
| M2759 | *MATα NUF2-3V5-IAA7:KanMX NDC80-3V5-IAA7:KanMX his3::pGPD1-OsTIR1:HIS3 leu2::pNUF2-nuf2(R133E)-3HA:LEU2 trp1::pNDC80-ndc80(ΔN-tail)-3HA:TRP1* | S6B |
| M2760 | *MATα NUF2-3V5-IAA7:KanMX NDC80-3V5-IAA7:KanMX his3::pGPD1-OsTIR1:HIS3 leu2::pNUF2-nuf2(R136E)-3HA:LEU2 trp1::pNDC80-ndc80(ΔN-tail)-3HA:TRP1* | S6B |
| M2763 | *MATα NUF2-3V5-IAA7:KanMX NDC80-3V5-IAA7:KanMX his3::pGPD1-OsTIR1:HIS3 leu2::pNUF2-nuf2(K92E)-3HA:LEU2 trp1::pNDC80-ndc80(ΔN-tail)-3HA:TRP1* | S6B |
| M2774 | *MATα NUF2-3V5-IAA7:KanMX NDC80-3V5-IAA7:KanMX his3::pGPD1-OsTIR1:HIS3 leu2::pNUF2-nuf2(F132A)-3HA:LEU2 trp1::pNDC80-ndc80(ΔN-tail)-3HA:TRP1* | 6B, S6B |
| M2792 | *MAT***a** *NUF2-3V5-IAA7:NatMX trp1::pGPD1-OsTIR1:TRP1 CDC20-IAA17:KanMX MTW1-mCherry:HygMX BUB1-GFP:KanMX leu2::pNUF2-NUF2-3HA::LEU2* | 2C |
| M2794 | *MAT***a** *NUF2-3V5-IAA7:NatMX trp1::pGPD1-OsTIR1:TRP1 CDC20-IAA17:KanMX MTW1-mCherry:HygMX BUB1-GFP:KanMX leu2::pNUF2-nuf2(S124D)-3HA::LEU2* | 2C |
| M2795 | *MAT***a** *NUF2-3V5-IAA7:NatMX trp1::pGPD1-OsTIR1:TRP1 CDC20-IAA17:KanMX MTW1-mCherry:HygMX BUB1-GFP:KanMX leu2::pNUF2-nuf2(F8A P9A)-3HA::LEU2* | 2C |
| M2801 | *MAT***a** *NUF2-3V5-IAA7:NatMX trp1::pGPD1-OsTIR1:TRP1 CDC20-IAA17:KanMX MTW1-mCherry:HygMX BUB1-GFP:KanMX leu2::pNUF2-nuf2(N128A)-3HA::LEU2* | 2C |
| M2841 | *MATα NUF2-3V5-IAA7:KanMX NDC80-3V5-IAA7:KanMX his3::pGPD1-OsTIR1:HIS3 leu2::pNUF2-nuf2(F132A)-3HA:LEU2 trp1::pNDC80-NDC80-3HA:TRP1* | 6B, S6A,B |
| M2850 | *MAT***a** *ura3::pGAL-MPS1-myc:URA3* | S6D |

| | | |
|---|---|---|
| M2867 | *MAT***a** *NUF2-3V5-IAA7:KanMX his3::pGPD1-OsTIR1:HIS3 DSN1-HIS-FLAG:URA3 leu2::pNUF2-NUF2-3HA:LEU2 ura3::pGAL-MPS1-myc:URA3 mad3Δ:::NatMX* | S6F |
| M2870 | *MAT***a** *NUF2-3V5-IAA7:KanMX his3::pGPD1-OsTIR1:HIS3 DSN1-HIS-FLAG:URA3 leu2::pNUF2-nuf2(F8A P9A)-3HA:LEU2 ura3::pGAL-MPS1-myc:URA3 mad3Δ:::NatMX* | S6F |
| M2933 | *MAT***a** *NUF2-3V5-IAA7:KanMX trp1::pGPD1-OsTIR1:TRP1 SPC110-mCherry:HygMX leu2::pNUF2-NUF2-3HA:LEU2* | 2B |
| M2935 | *MAT***a** *NUF2-3V5-IAA7:KanMX trp1::pGPD1-OsTIR1:TRP1 SPC110-mCherry:HygMX leu2::pNUF2-nuf2(S124D)-3HA:LEU2* | 2B |
| M2936 | *MAT***a** *NUF2-3V5-IAA7:KanMX trp1::pGPD1-OsTIR1:TRP1 SPC110-mCherry:HygMX leu2::pNUF2-nuf2(F8A P9A)-3HA:LEU2* | 2B |
| M2945 | *MAT***a** *NUF2-3V5-IAA7:KanMX trp1::pGPD1-OsTIR1:TRP1 SPC110-mCherry:HygMX leu2::pNUF2-nuf2(N128A)-3HA:LEU2* | 2B |
| M2998 | *MATα NUF2-3V5-IAA7:KanMX NDC80-3V5-IAA7:KanMX his3::pGPD1-OsTIR1:HIS3 leu2::pNUF2-NUF2-3HA:LEU2 trp1::pNDC80-NDC80-3HA:TRP1* | 6B, S6A,B |
| M2999 | *MATα NUF2-3V5-IAA7:KanMX NDC80-3V5-IAA7:KanMX his3::pGPD1-OsTIR1:HIS3 leu2::pNUF2-NUF2-3HA:LEU2 trp1::pNDC80-ndc80(ΔN-tail)-3HA:TRP1* | 6B, S6B |
| M3441 | *MAT***a** *NUF2-3V5-IAA7:KanMX trp1::pGPD1-OsTIR1:TRP1 DSN1-HIS-FLAG:URA3 leu2::pNUF2-nuf2(D39K)-3HA:LEU2* | S1D |
| M3442 | *MAT***a** *NUF2-3V5-IAA7:KanMX trp1::pGPD1-OsTIR1:TRP1 DSN1-HIS-FLAG:URA3 leu2::pNUF2-nuf2(E134K)-3HA:LEU2* | S1D |
| M3443 | *MAT***a** *NUF2-3V5-IAA7:KanMX trp1::pGPD1-OsTIR1:TRP1 DSN1-HIS-FLAG:URA3 leu2::pNUF2-nuf2(E15K)-3HA:LEU2* | S1D |
| M3444 | *MAT***a** *NUF2-3V5-IAA7:KanMX trp1::pGPD1-OsTIR1:TRP1 DSN1-HIS-FLAG:URA3 leu2::pNUF2-nuf2(E58K)-3HA:LEU2* | S1D |
| M3494 | *MAT***a** *NUF2-3V5-IAA7:KanMX his3::pGPD1-OsTIR1:HIS3 DSN1-HIS-FLAG:URA3 MPS1-3V5:HygMX leu2::pNUF2-NUF2-3HA:LEU2* | 4A |
| M3498 | *MAT***a** *NUF2-3V5-IAA7:KanMX his3::pGPD1-OsTIR1:HIS3 DSN1-HIS-FLAG:URA3 MPS1-3V5:HygMX leu2::pNUF2-nuf2(S124D)-3HA:LEU2* | 4A |
| M3500 | *MAT***a** *NUF2-3V5-IAA7:KanMX his3::pGPD1-OsTIR1:HIS3 DSN1-HIS-FLAG:URA3 MPS1-3V5:HygMX leu2::pNUF2-nuf2(F8A P9A)-3HA:LEU2* | 4A |
| M3504 | *MAT***a** *NUF2-3V5-IAA7:KanMX his3::pGPD1-OsTIR1:HIS3 DSN1-HIS-FLAG:URA3 MPS1-3V5:HygMX leu2::pNUF2-nuf2(N128A)-3HA:LEU2* | 4A |
| M3507 | *MAT***a** *NUF2-3V5-IAA7:KanMX trp1::pGPD1-OsTIR1:TRP1 DSN1-HIS-FLAG:URA3 leu2::pNUF2-nuf2(E15K E134K)-3HA:LEU2* | S1D |
| M3586 | *MAT***a** *NUF2-3V5-IAA7:KanMX trp1::pGPD1-OsTIR1:TRP1 DSN1-HIS-FLAG:URA3 leu2::pNUF2-NUF2-3HA:LEU2 his3::pGAL10-MPS1:HIS3* | 6C, S6G |
| M3587 | *MAT***a** *NUF2-3V5-IAA7:KanMX trp1::pGPD1-OsTIR1:TRP1 DSN1-HIS-FLAG:URA3 leu2::pNUF2-nuf2(F8A P9A)-3HA:LEU2 his3::pGAL10-MPS1:HIS3* | 6C, S6G,H, I |
| M3588 | *MAT***a** *NUF2-3V5-IAA7:KanMX trp1::pGPD1-OsTIR1:TRP1 DSN1-HIS-FLAG:URA3 leu2::pNUF2-nuf2(N128A)-3HA:LEU2 his3::pGAL10-MPS1:HIS3* | 6C,E,G |
| M3750 | *MAT***a** *NUF2-3V5-IAA7:KanMX trp1::pGPD1-OsTIR1:TRP1 DSN1-HIS-FLAG:URA3 leu2::pNUF2-nuf2(I10A L11A)-3HA:LEU2* | 5B, S1D |
| M3751 | *MAT***a** *NUF2-3V5-IAA7:KanMX trp1::pGPD1-OsTIR1:TRP1 DSN1-HIS-FLAG:URA3 leu2::pNUF2-nuf2(I10A S124A)-3HA:LEU2* | S1D |

| | | |
|---|---|---|
| M3752 | *MAT***a** *NUF2-3V5-IAA7:KanMX trp1::pGPD1-OsTIR1:TRP1 DSN1-HIS-FLAG:URA3 leu2::pNUF2-nuf2(L11A S124A)-3HA:LEU2* | S1D |
| M3753 | *MAT***a** *NUF2-3V5-IAA7:KanMX his3::pGPD1-OsTIR1:HIS3 SPC110-mCherry:HygMX BUB1-GFP:KanMX pMET-CDC20:TRP1 leu2::pNUF2-NUF2-3HA:LEU2* | S3A |
| M3757 | *MAT***a** *NUF2-3V5-IAA7:KanMX his3::pGPD1-OsTIR1:HIS3 SPC110-mCherry:HygMX BUB1-GFP:KanMX pMET-CDC20:TRP1 leu2::pNUF2-nuf2(S124D)-3HA:LEU2* | S3A |
| M3874 | *MAT***a** *NUF2-3V5-IAA7:NatMX trp1::pGPD1-OsTIR1:TRP1 CEN III-lacO128:TRP1 his3::pCUP1-GFP12-lacI12:HIS3 SPC110-mCherry:HygMX CDC20-IAA17:KanMX leu2::pNUF2-NUF2-3HA:LEU2* | S3B,C |
| M3878 | *MAT***a** *NUF2-3V5-IAA7:NatMX trp1::pGPD1-OsTIR1:TRP1 CEN III-lacO128:TRP1 his3::pCUP1-GFP12-lacI12:HIS3 SPC110-mCherry:HygMX CDC20-IAA17:KanMX leu2::pNUF2-nuf2(S124D-3HA:LEU2* | S3B,C |
| M3996 | *MAT***a** *NUF2-3V5-IAA7:KanMX trp1::pGPD1-OsTIR1:TRP1 DSN1-HIS-FLAG:URA3 leu2::pNUF2-nuf2(F8D)-3HA:LEU2* | S1D, S2 |
| M4055 | *MAT***a** *NUF2-3V5-IAA7:KanMX trp1::pGPD1-OsTIR1:TRP1 DSN1-HIS-FLAG:URA3 leu2::pNUF2-nuf2(N128A)-3HA:LEU2 his3::pGAL10-mps1(Δ151-200):HIS3* | S6H |
| M4056 | *MAT***a** *NUF2-3V5-IAA7:KanMX trp1::pGPD1-OsTIR1:TRP1 DSN1-HIS-FLAG:URA3 leu2::pNUF2-nuf2(N128A)-3HA:LEU2 his3::pGAL10-mps1(Δ201-300):HIS3* | S6H |
| M4116 | *MAT***a** *TOR1-1 fpr1Δ::NatMX NDC80-FKBP12:His3MX MPS1-FRB:KanMX mad3Δ::HygMX* | 7B |
| M4171 | *MAT***a** *NUF2-3V5-IAA7:KanMX trp1::pGPD1-OsTIR1:TRP1 DSN1-HIS-FLAG:URA3 leu2::pNUF2-NUF2-3HA:LEU2 his3::pGAL10-IPL1::HIS3* | S6G |
| M4172 | *MAT***a** *NUF2-3V5-IAA7:KanMX trp1::pGPD1-OsTIR1:TRP1 DSN1-HIS-FLAG:URA3 leu2::pNUF2-nuf2(F8A P9A)-3HA:LEU2 his3::pGAL10-IPL1::HIS3* | S6G |
| M4173 | *MAT***a** *NUF2-3V5-IAA7:KanMX trp1::pGPD1-OsTIR1:TRP1 DSN1-HIS-FLAG:URA3 leu2::pNUF2-nuf2(N128A)-3HA:LEU2 his3::pGAL10-IPL1::HIS3* | S6G |
| M4182 | *MAT***a** *NDC80-3V5-IAA7:KanMX leu2::pGPD1-OsTIR1:LEU2* | 2A |
| M4245 | *MAT***a** *NUF2-3V5-IAA7:KanMX ura3::pGPD1-OsTIR1:URA3 CEN IV-lacO128:TRP1 his3::pCUP1-GFP12-LacI12:HIS3 pMET-CDC20:TRP1 leu2::pNUF2-NUF2-3HA:LEU2* | 3B |
| M4249 | *MAT***a** *NUF2-3V5-IAA7:KanMX ura3::pGPD1-OsTIR1:URA3 CEN IV-lacO128:TRP1 his3::pCUP1-GFP12-LacI12:HIS3 pMET-CDC20:TRP1 leu2::pNUF2-nuf2(S124D)-3HA:LEU2* | 3B |
| M4251 | *MAT***a** *NUF2-3V5-IAA7:KanMX ura3::pGPD1-OsTIR1:URA3 CEN IV-lacO128:TRP1 his3::pCUP1-GFP12-LacI12:HIS3 pMET-CDC20:TRP1 leu2::pNUF2-nuf2(F8A P9A)-3HA:LEU2* | 3B |
| M4253 | *MAT***a** *NUF2-3V5-IAA7:KanMX ura3::pGPD1-OsTIR1:URA3 CEN IV-lacO128:TRP1 his3::pCUP1-GFP12-LacI12:HIS3 pMET-CDC20:TRP1 leu2::pNUF2-nuf2(N128A)-3HA:LEU2* | 3B |
| M4407 | *MAT***a** *mps1-1 his3::pMPS1-MPS1:HIS3* | 4C, S4A |
| M4409 | *MAT***a** *mps1-1 his3::pMPS1-mps1(Δ151-157):HIS3* | S4A |
| M4410 | *MAT***a** *mps1-1 his3::pMPS1-mps1(Δ165-171):HIS3* | S4A |

| | | |
|---|---|---|
| M4411 | *MAT***a** *mps1-1 his3::pMPS1-mps1(D580A):HIS3* | S4A |
| M4412 | *MAT***a** *mps1-1 his3::pMPS1-mps1(∆151-200):HIS3* | 4C, S4A |
| M4413 | *MAT***a** *mps1-1 his3::pMPS1-mps1(∆201-300):HIS3* | 4C, S4A |
| M4414 | *MAT***a** *mps1-1 his3::pMPS1-mps1(L165E R170E):HIS3* | S4A |
| M4415 | *MAT***a** *mps1-1 his3::pMPS1-mps1(R151E R152E K154E R155E):HIS3* | S4A |
| M4416 | *MAT***a** *mps1-1 his3::pMPS1-mps1(K169E R170E):HIS3* | S4A |
| M4417 | *MAT***a** *mps1-1 his3::pMPS1-mps1(∆4-50):HIS3* | S4A |
| M4418 | *MAT***a** *mps1-1 his3::pMPS1-mps1(∆151-157, ∆165-171):HIS3* | S4A |
| M4420 | *MAT***a** *mps1-1 his3::pMPS1-mps1(R151E R152E K154E R155E L169E R170E):HIS3* | S4A |
| M4465 | *MAT***a** *NUF2-3V5-IAA7:NatMX his3::pGPD1-OsTIR1:HIS3 DSN1-HIS-FLAG:URA3 trp1::256lacO:TRP1 dad1-1:KanMX leu2::pNUF2-NUF2-3HA:LEU2* | 5B, S5 |
| M4469 | *MAT***a** *NUF2-3V5-IAA7:KanMX ura3::pGPD1-OsTIR1:URA3 CEN IV-lacO128:TRP1 his3::pCUP1-GFP12-LacI12:HIS3 SPC110-mCherry:HygMX leu2::pNUF2-NUF2-3HA:LEU2* | 3C, S3D |
| M4475 | *MAT***a** *NUF2-3V5-IAA7:KanMX ura3::pGPD1-OsTIR1:URA3 CEN IV-lacO128:TRP1 his3::pCUP1-GFP12-LacI12:HIS3 SPC110-mCherry:HygMX leu2::pNUF2-nuf2(F8A P9A)-3HA:LEU2* | 3C, S3D |
| M4479 | *MAT***a** *NUF2-3V5-IAA7:KanMX ura3::pGPD1-OsTIR1:URA3 CEN IV-lacO128:TRP1 his3::pCUP1-GFP12-LacI12:HIS3 SPC110-mCherry:HygMX leu2::pNUF2-nuf2(N128A)-3HA:LEU2* | 3C |
| M4618 | *MAT***a** *his3::pGAL10-MPS1:HIS3* | S6D |
| M4619 | *MAT***a** *NUF2-3V5-IAA7:KanMX trp1::pGPD1-OsTIR1:TRP1 DSN1-HIS-FLAG:URA3 leu2::pNUF2-nuf2(S124D)-3HA:LEU2 his3::pGAL10-MPS1:HIS3* | 6C |
| M4640 | *MAT***a** *NUF2-3V5-IAA7:KanMX trp1::pGPD1-OsTIR1:TRP1 DSN1-HIS-FLAG:URA3 leu2::pNUF2-nuf2(N128A)-3HA:LEU2 his3::pGAL10-mps1(L165E R170E):HIS3* | S6I |
| M4641 | *MAT***a** *NUF2-3V5-IAA7:KanMX trp1::pGPD1-OsTIR1:TRP1 DSN1-HIS-FLAG:URA3 leu2::pNUF2-nuf2(N128A)-3HA:LEU2 his3::pGAL10-mps1(R151E R152E K154E R155E):HIS3* | S6I |
| M4642 | *MAT***a** *NUF2-3V5-IAA7:KanMX trp1::pGPD1-OsTIR1:TRP1 DSN1-HIS-FLAG:URA3 leu2::pNUF2-nuf2(N128A)-3HA:LEU2 his3::pGAL10-mps1(K169E R170E):HIS3* | S6I |
| M4643 | *MAT***a** *NUF2-3V5-IAA7:KanMX trp1::pGPD1-OsTIR1:TRP1 DSN1-HIS-FLAG:URA3 leu2::pNUF2-nuf2(N128A)-3HA:LEU2 his3::pGAL10-mps1(∆151-157 ∆165-171):HIS3* | S6I |
| M4645 | *MAT***a** *NUF2-3V5-IAA7:KanMX trp1::pGPD1-OsTIR1:TRP1 DSN1-HIS-FLAG:URA3 leu2::pNUF2-nuf2(N128A)-3HA:LEU2 his3::pGAL10-mps1(R151E R152E K154E R155E K169E R170E):HIS3* | S6I |
| M4710 | *MAT***a** *mps1-1 CEN IV-lacO128:TRP1 his3::pCUP1-GFP12-LacI12:HIS3 SPC110-mCherry:HygMX ura3::pMPS1-mps1(R155E L165E R170E):URA3* | 4D, S4B,C |
| M4714 | *MAT***a** *mps1-1 CEN IV::lacO128:TRP1 his3::pCUP1-GFP12-LacI12:HIS3 SPC110-mCherry:HygMX ura3::pMPS1-MPS1:URA3* | 4D, S4B,C |
| M4841 | *MAT***a** *TOR1-1 fpr1∆::NatMX NDC80-FKBP12:His3MX MPS1-FRB:KanMX mad3∆::HygMX NUF2-3V5-IAA7:KanMX trp1::pGPD1-OsTIR1:TRP1* | 7B |

| | | |
|---|---|---|
| M4908 | *MAT***a** *TOR1-1 fpr1Δ::NatMX NDC80-FKBP12:His3MX MPS1-FRB:KanMX mad3Δ::HygMX NUF2-3V5-IAA7:KanMX trp1::pGPD1-OsTIR1:TRP1 leu2::pNUF2-NUF2-3HA:LEU2* | 7B |
| M4912 | *MAT***a** *TOR1-1 fpr1Δ::NatMX NDC80-FKBP12:His3MX MPS1-FRB:KanMX mad3Δ::HygMX NUF2-3V5-IAA7:KanMX trp1::pGPD1-OsTIR1:TRP1 leu2::pNUF2-nuf2(N128A)-3HA:LEU2* | 7B |
| M4924 | *MAT***a** *NUF2-3V5-IAA7:NatMX his3::pGPD1-OsTIR1:HIS3 DSN1-HIS-FLAG:URA3 trp1-1:256lacO:TRP1 dad1-1:KanMX leu2::pNUF2-nuf2(F8A)-3HA:LEU2* | S5 |
| M4926 | *MAT***a** *NUF2-3V5-IAA7:NatMX his3::pGPD1-OsTIR1:HIS3 DSN1-HIS-FLAG:URA3 trp1::256lacO:TRP1 dad1-1:KanMX leu2::pNUF2-nuf2(P9A)-3HA:LEU2* | 5B |
| M4928 | *MAT***a** *NUF2-3V5-IAA7:NatMX his3::pGPD1-OsTIR1:HIS3 DSN1-HIS-FLAG:URA3 trp1::256lacO:TRP1 dad1-1:KanMX leu2::pNUF2-nuf2(I10A L11A)-3HA:LEU2* | 5B |
| M4946 | *MAT***a** *NUF2-3V5-IAA7:KanMX trp1::pGPD1-OsTIR1:TRP1 DSN1-HIS-FLAG:URA3 leu2::pNUF2-nuf2(N128A)-3HA:LEU2 his3::pGAL10-mps1(R155E L165E R170E):HIS3* | 6E, S6H,I |
| M4947 | *MAT***a** *mps1-1 CEN IV::lacO128:TRP1 his3::pCUP1-GFP12-LacI12:HIS3 SPC110-mCherry:HygMX ura3::pMPS1-mps1(R155E L165E R170E)::URA3* | 4D, S4B,C |
| M4948 | *MAT***a** *mps1-1 CEN IV::lacO128:TRP1 his3::pCUP1-GFP12-LacI12:HIS3 SPC110-mCherry:HygMX ura3::pMPS1-mps1(R155E L165E R170E)::URA3* | 4D, S4C |
| M4949 | *MAT***a** *mps1-1 his3::pMPS1-mps1(R155E L165E R170E):HIS3* | 4C, S4A |
| M4955 | *MAT***a** *TOR1-1 fpr1Δ::NatMX MPS1-FRB:KanMX mad3Δ::HygMX NUF2-3V5-IAA7:KanMX trp1::pGPD1-OsTIR1:TRP1 leu2::pNUF2-nuf2(N128A)-3HA:LEU2* | 7B |
| M4985 | *MAT***a** *trp1::pGPD1-OsTIR1:TRP1 CDC20-AID:KanMX Spc110-mCherry:HphMX NUF2-3V5-IAA7:KanMX Ask1-YFP:HIS3 leu2::pNUF2-NUF2-3HA:LEU2* | 5A |
| M4987 | *MAT***a** *trp1::pGPD1-OsTIR1:TRP1 CDC20-AID:KanMX Spc110-mCherry:HphMX NUF2-3V5-IAA7:KanMX Ask1-YFP:HIS3 leu2::pNUF2-nuf2(F8A P9A)-3HA:LEU2* | 5A |

**Table S2. Plasmids and primers used for strain construction.**

| **Plasmid** | **Primers used to generate plasmids (5' to 3'):** | |
|---|---|---|
| **pM270** (*pNDC80-NDC80-3HA*, i.e. *NDC80*$^{WT}$) | SB4732 | GATCGATCgggcccGGTCTCTGTAGGGTCAATAG |
| | SB4443 | GATCGATCtctagaTCAGCACTGAGCAGCGTAATCTGGAACG |
| **pM272** (*pNDC80-ndc80(K204E)-3HA*, i.e. *ndc80*$^{K204E}$) | SB1961 | CGTATGGAATGATATAGTCAC |
| | SB4752 | CCTACAGCCGAAATTTGTGATTcATTTATTGACTCTAAAAACG |
| **pM273** (*pNDC80-ndc80(K122E K204E)-3HA*, i.e. *ndc80*$^{K122E\ K204E}$) | SB4751 | GATCCAAGGCCACTAAGAGACgAAAACTTCCAAAGCGCTATTCAAG |
| | SB4752 | CCTACAGCCGAAATTTGTGATTcATTTATTGACTCTAAAAACG |
| **pM604** (*pNUF2-NUF2-3HA*, i.e. *NUF2*$^{WT}$) | oMM196 | taccGGGCCCTCAACGCCTTCTTGAATAATTC |
| | oMM197 | ctagtttctagaactatcagcactgagcagcg |
| **pM633** (*pNUF2-nuf2(S124A)-3HA*, i.e. *nuf2*$^{S124A}$) | oMM199 | GGTATATTATCCCCATCTACGTCCTCGTATTGCTTTAGC |
| | oMM200 | CAACGGACACAGCGCTTACTGGCTGCTGTGGTGAATTACGCTCG |
| **pM638** (*pNUF2-nuf2(S124D)-3HA*, i.e. *nuf2*$^{S124D}$) | oMM199 | GGTATATTATCCCCATCTACGTCCTCGTATTGCTTTAGC |
| | oMM242 | CAACGGACACAGCGCTTACTGGATGCTGTGGTGAATTACGCTCG |
| **pM639** (*pNUF2-nuf2(F8A P9A)-3HA*, i.e. *nuf2*$^{F8A\ P9A}$) | oMM199 | GGTATATTATCCCCATCTACGTCCTCGTATTGCTTTAGC |
| | oMM243 | AGTAGGAATCAAGATGTGGCTGCAATTTTGGATCTACAGGAACTAG |
| **pM640** (*pNUF2-nuf2(S2A R3A N4A Q5A)-3HA*, i.e. *nuf2*$^{S2A\ R3A\ N4A\ Q5A}$) | oMM199 | GGTATATTATCCCCATCTACGTCCTCGTATTGCTTTAGC |
| | oMM244 | CATCCCTTGAGCAAAATGGCTGCAGCCGCTGATGTGTTCCCCATTTTGGATC |
| **pM641** (*pNUF2-nuf2(D6A V7A I10A L11A)-3HA*, i.e. *nuf2*$^{D6A\ V7A\ I10A\ L11A}$) | oMM199 | GGTATATTATCCCCATCTACGTCCTCGTATTGCTTTAGC |
| | oMM245 | AAAATGAGTAGGAATCAAGCTGCATTCCCCGCCGCTGATCTACAGGAACTAGTTATATG |
| **pM642** (*pNUF2-nuf2(Δ2-11)-3HA*, i.e. *nuf2*$^{\Delta 2\text{-}11}$) | oMM199 | GGTATATTATCCCCATCTACGTCCTCGTATTGCTTTAGC |
| | oMM246 | CTCCAGCATCCCTTGAGCAAAATGGATCTACAGGAACTAGTTATATG |
| **pM643** (*pNUF2-nuf2(Δ6-7)-3HA*, i.e. *nuf2*$^{\Delta 6\text{-}7}$) | oMM199 | GGTATATTATCCCCATCTACGTCCTCGTATTGCTTTAGC |
| | oMM247 | TTGAGCAAAATGAGTAGGAATCAATTCCCCATTTTGGATCTACAGG |

| | | |
|---|---|---|
| **pM644** (*pNUF2-nuf2(Δ8-11)-3HA*, i.e. *nuf2*$^{\Delta 8\text{-}11}$) | oMM199 | GGTATATTATCCCCATCTACGTCCTC GTATTGCTTTAGC |
| | oMM248 | AAAATGAGTAGGAATCAAGATGTGGA TCTACAGGAACTAGTTATATG |
| **pM645** (*pNUF2-nuf2(Δ2-5)-3HA*, i.e. *nuf2*$^{\Delta 2\text{-}5}$) | oMM199 | GGTATATTATCCCCATCTACGTCCTC GTATTGCTTTAGC |
| | oMM249 | CTCCAGCATCCCTTGAGCAAAATGG ATGTGTTCCCCATTTTGGATC |
| **pM672** (*pNUF2-nuf2(F8A)-3HA*, i.e. *nuf2*$^{F8A}$) | oMM199 | GGTATATTATCCCCATCTACGTCCTC GTATTGCTTTAGC |
| | oMM280 | AGTAGGAATCAAGATGTGGCTCCCAT TTTGGATCTACAGGAACTAG |
| **pM673** (*pNUF2-nuf2(P9A)-3HA*, i.e. *nuf2*$^{P9A}$) | oMM199 | GGTATATTATCCCCATCTACGTCCTC GTATTGCTTTAGC |
| | oMM281 | AGTAGGAATCAAGATGTGTTCGCAAT TTTGGATCTACAGGAACTAG |
| **pM674** (*pNUF2-nuf2(K113E)-3HA*, i.e. *nuf2*$^{K113E}$) | oMM285 | CTGAGGAGAAAGGCTCCAGCATCCC TTGAGCAAAATG |
| | oMM282 | TCCGTTGGGCTTCGGGTTCGTACAA ATCTGTCATATTGAAATC |
| **pM675** (*pNUF2-nuf2(R118E)-3HA*, i.e. *nuf2*$^{R118E}$) | oMM285 | CTGAGGAGAAAGGCTCCAGCATCCC TTGAGCAAAATG |
| | oMM283 | TCAGTAAGCGCTGTGTTTCTTGGGC TTCGGGCTTGTACAAATC |
| **pM699** (*pNUF2-nuf2(I10A)-3HA*, i.e. *nuf2*$^{I10A}$) | oMM314 | AGTAGGAATCAAGATGTGTTCCCCG CTTTGGATCTACAGGAACTAG |
| | oMM199 | GGTATATTATCCCCATCTACGTCCTC GTATTGCTTTAGC |
| **pM700** (*pNUF2-nuf2(L11A)-3HA*, i.e. *nuf2*$^{L11A}$) | oMM315 | AGTAGGAATCAAGATGTGTTCCCCAT TGCTGATCTACAGGAACTAG |
| | oMM199 | GGTATATTATCCCCATCTACGTCCTC GTATTGCTTTAGC |
| **pM714** (*pNUF2-nuf2(R121E)-3HA*, i.e. *nuf2*$^{R121E}$) | oMM285 | CTGAGGAGAAAGGCTCCAGCATCCC TTGAGCAAAATG |
| | oMM318 | CACAGCACTCAGTAATTCCTGTGTCC GTTGGGCTTCGGGCTT |
| **pM715** (*pNUF2-nuf2(R131E)-3HA*, i.e. *nuf2*$^{R131E}$) | oMM285 | CTGAGGAGAAAGGCTCCAGCATCCC TTGAGCAAAATG |
| | oMM319 | TCGTTCCTCCCTAAATTCAGCGTAAT TCACCACAGCACTCAG |
| **pM716** (*pNUF2-nuf2(R133E)-3HA*, i.e. *nuf2*$^{R133E}$) | oMM285 | CTGAGGAGAAAGGCTCCAGCATCCC TTGAGCAAAATG |
| | oMM320 | CGAACATTCGTTCCTCTTCAAAACGA GCGTAATTCACCACAG |
| **pM717** (*pNUF2-nuf2(R136E)-3HA*, i.e. *nuf2*$^{R136E}$) | oMM285 | CTGAGGAGAAAGGCTCCAGCATCCC TTGAGCAAAATG |
| | oMM321 | ATTACAGTCGAACATTTCTTCCTCCC TAAAACGAGCGTAATT |

| | | |
|---|---|---|
| **pM718** (*pNUF2-nuf2(R35E)-3HA*, i.e. $nuf2^{R35E}$) | oMM329 | CCACACAGGAAAATATCTCTGAACCCACCTCAGACTACATGG |
| | oMM199 | GGTATATTATCCCCATCTACGTCCTCGTATTGCTTTAGC |
| **pM719** (*pNUF2-nuf2(K49E)-3HA*, i.e. $nuf2^{K49E}$) | oMM332 | ACTACATGGTAACCCTTTACGAACAAATCATCGAGAACTTCAT |
| | oMM199 | GGTATATTATCCCCATCTACGTCCTCGTATTGCTTTAGC |
| **pM720** (*pNUF2-nuf2(K92E)-3HA*, i.e. $nuf2^{K92E}$) | oMM330 | TAAATGTTTTGGTATTGAACGAAATCTGCTTTAAGTTCTTTG |
| | oMM199 | GGTATATTATCCCCATCTACGTCCTCGTATTGCTTTAGC |
| **pM721** (*pNUF2-nuf2(K96E)-3HA*, i.e. $nuf2^{K96E}$) | oMM331 | TATTGAACAAAATCTGCTTTGAATTCTTTGAGAACATAGGTG |
| | oMM199 | GGTATATTATCCCCATCTACGTCCTCGTATTGCTTTAGC |
| **pM733** (*pNUF2-nuf2(F8A P9A L11A)-3HA*, i.e. $nuf2^{F8A\ P9A\ L11A}$) | oMM337 | AGTAGGAATCAAGATGTGGCTGCAATTGCTGATCTACAGGAACTAG |
| | oMM199 | GGTATATTATCCCCATCTACGTCCTCGTATTGCTTTAGC |
| **pM734** (*pNUF2-nuf2(R118E R133E)-3HA*, i.e. $nuf2^{R118E\ R133E}$) | oMM285 | CTGAGGAGAAAGGCTCCAGCATCCCTTGAGCAAAATG |
| | oMM320 | CGAACATTCGTTCCTCTTCAAAACGAGCGTAATTCACCACAG |
| **pM735** (*pNUF2-nuf2(R118E R121E)-3HA*, i.e. $nuf2^{R118E\ R121E}$) | oMM285 | CTGAGGAGAAAGGCTCCAGCATCCCTTGAGCAAAATG |
| | oMM318 | CACAGCACTCAGTAATTCCTGTGTCCGTTGGGCTTCGGGCTT |
| **pM750** (*pNUF2-nuf2(K113E R118E)-3HA*, i.e. $nuf2^{K113E\ R118E}$) | oMM285 | CTGAGGAGAAAGGCTCCAGCATCCCTTGAGCAAAATG |
| | oMM338 | CACAGCACTCAGTAAGCGCTGTGTTTCTTGGGCTTCGGGTTCGTACAAATCTGTCATATTGAAATC |
| **pM777** (*pNUF2-nuf2(D105K)-3HA*, i.e. $nuf2^{D105K}$) | oMM285 | CTGAGGAGAAAGGCTCCAGCATCCCTTGAGCAAAATG |
| | oMM368 | GTACAAATCTGTCATATTGAATTTTTGAACACCTATGTTCTCAAAG |
| **pM778** (*pNUF2-nuf2(N128A)-3HA*, i.e. $nuf2^{N128A}$) | oMM285 | CTGAGGAGAAAGGCTCCAGCATCCCTTGAGCAAAATG |
| | oMM369 | CCTCCCTAAAACGAGCGTATGCCACCACAGCACTCAGTAAGC |
| **pM1342** (*pNDC80-ndc80(ΔN-term)-3HA*, i.e. $ndc80^{\Delta N\text{-}tail}$) | oMM223 | GAGAGGTAGAATCGTCCCTG |
| | oMM376 | CTCCTCTTGAATAGCGCTTTGGAAGTTTTTGTCTCTTAGTGGCCTTGGATCTCTATTcatttatagaaacgggtatc |
| **pM1355** (*pNUF2-nuf2(A125D)-3HA*, i.e. $nuf2^{A125D}$) | oMM285 | CTGAGGAGAAAGGCTCCAGCATCCCTTGAGCAAAATG |
| | oMM395 | AAACGAGCGTAATTCACCACATCACTCAGTAAGCGCTGTGTCCGTTG |

| | | |
|---|---|---|
| **pM1359** (*pNUF2-nuf2(D110K)-3HA*, i.e. *nuf2*$^{D110K}$) | oMM285 | CTGAGGAGAAAGGCTCCAGCATCCCTTGAGCAAAATG |
| | oMM393 | GGGCTTCGGGCTTGTACAATTTTGTCATATTGAAATCTTGAAC |
| **pM1360** (*pNUF2-nuf2(F132A)-3HA*, i.e. *nuf2*$^{F132A}$) | oMM285 | CTGAGGAGAAAGGCTCCAGCATCCCTTGAGCAAAATG |
| | oMM396 | CGAACATTCGTTCCTCCCTAGCACGAGCGTAATTCACCACAGCAC |
| **pM1475** (*pNUF2-nuf2(D39K)-3HA*, i.e. *nuf2*$^{D39K}$) | oMM543 | GGAAAATATCTCTAGGCCCACCTCAAAATACATGGTAACCCTTTACAAAC |
| | oMM199 | GGTATATTATCCCCATCTACGTCCTCGTATTGCTTTAGC |
| **pM1476** (*pNUF2-nuf2(E134K)-3HA*, i.e. *nuf2*$^{E134K}$) | oMM545 | AGAATTACAGTCGAACATTCGTTCTTTCCTAAAACGAGCGTAATTC |
| | oMM199 | GGTATATTATCCCCATCTACGTCCTCGTATTGCTTTAGC |
| **pM1477** (*pNUF2-nuf2(E15K)-3HA*, i.e. *nuf2*$^{E15K}$) | oMM542 | TGTGTTCCCCATTTTGGATCTACAGAAACTAGTTATATGTTTGCAAAGC |
| | oMM199 | GGTATATTATCCCCATCTACGTCCTCGTATTGCTTTAGC |
| **pM1478** (*pNUF2-nuf2(E58K)-3HA*, i.e. *nuf2*$^{E58K}$) | oMM544 | ATCGAGAACTTCATGGGTATTTCTGTAAAATCGTTGCTGAATAGTAGTAAC |
| | oMM199 | GGTATATTATCCCCATCTACGTCCTCGTATTGCTTTAGC |
| **pM1485** (*pNUF2-nuf2(E15K E134K)-3HA*, i.e. *nuf2*$^{E15K\ E134K}$) | oMM542 | TGTGTTCCCCATTTTGGATCTACAGAAACTAGTTATATGTTTGCAAAGC |
| | oMM199 | GGTATATTATCCCCATCTACGTCCTCGTATTGCTTTAGC |
| **pM1489** (*pGAL10-MPS1*, i.e. *MPS1*$^{WT}$) | oMM565 | gcaaggcaggtggtcgacggtatcgataagcttgatatcgATGTCAACAAACTCATTCCATG |
| | oMM566 | ttgcaggtgtctagaactagtggatcccccgggctgcaggCTAAATTTTGTAATCTGCAAATTTCC |
| **pM1490** (*pGAL10-IPL1*, i.e. *IPL1*$^{WT}$) | oMM567 | gcaaggcaggtggtcgacggtatcgataagcttgatatcgATGCAACGCAATAGTTTAGTAAATATC |
| | oMM568 | ttgcaggtgtctagaactagtggatcccccgggctgcaggCTATAACCGCTTATTTTCCC |
| **pM1526** (*pNUF2-nuf2(I10A L11A)-3HA*, i.e. *nuf2*$^{I10A\ L11A}$) | oMM660 | AGTAGGAATCAAGATGTGTTCCCCGCTGCTGATCTACAGGAACTAG |
| | oMM199 | GGTATATTATCCCCATCTACGTCCTCGTATTGCTTTAGC |
| **pM1527** (*pNUF2-nuf2(I10A S124A)-3HA*, i.e. *nuf2*$^{I10A\ S124A}$) | oMM314 | AGTAGGAATCAAGATGTGTTCCCCGCTTTGGATCTACAGGAACTAG |
| | oMM199 | GGTATATTATCCCCATCTACGTCCTCGTATTGCTTTAGC |

| | | |
|---|---|---|
| **pM1528** (*pNUF2-nuf2(L11A S124A)-3HA*, i.e. *nuf2*$^{L11A\ S124A}$) | oMM315 | AGTAGGAATCAAGATGTGTTCCCCATTGCTGATCTACAGGAACTAG |
| | oMM199 | GGTATATTATCCCCATCTACGTCCTCGTATTGCTTTAGC |
| **pM1550** (*pNUF2-nuf2(F8D)-3HA*, i.e. *nuf2*$^{F8D}$) | oMM696 | AGTAGGAATCAAGATGTGGATCCCATTTTGGATCTACAGGAACTAG |
| | oMM199 | GGTATATTATCCCCATCTACGTCCTCGTATTGCTTTAGC |
| **pM1578** (*pGAL10-mps1(Δ151-200)*, i.e. *mps1*$^{\Delta 151-200}$) | oMM713 | gcgacaaaatatgaaagaagatattacggcaaagtatgctgaaGAGGATTCTCACCAAACAAAC |
| | oMM570 | GTTGAAGGAGATTATCAGCG |
| **pM1579** (*pGAL10-mps1(Δ201-300),* i.e. *mps1*$^{\Delta 201-300}$) | oMM714 | ccagccaataaatgcaagggagacagtggaattaccacttCCCAGGCGAAAAGTTTCTAC |
| | oMM570 | GTTGAAGGAGATTATCAGCG |
| **pM1595** (*pMPS1-MPS1*, i.e. *MPS1*$^{WT}$) *HIS3* SIV | oMM761 | taccGGGCCCTGTTATCACAACAAATGGTGATTCTGG |
| | oMM762 | atagtaccGTCGACGTTTGTTTTGAGATCATCCAGTTCTTG |
| **pM1608** (*pMPS1-mps1(Δ151-157),* i.e. *mps1*$^{\Delta 151-157}$) *HIS3* SIV | oMM757 | gcgacaaaatatgaaagaagatattacggcaaagtatgctgaaATATCCAATAGGACAACGAAGC |
| | oMM570 | GTTGAAGGAGATTATCAGCG |
| **pM1609** (*pMPS1-mps1(Δ165-171),* i.e. *mps1*$^{\Delta 165-171}$) *HIS3* SIV | oMM758 | aaggagaagtaagagatttttaatatccaataggacaacg GAAAGGTATCTTAAAAATCATTGCATTTGGTATAGCAAACGCGG aagATGACTTTGACAAATATCTTTGATGAG |
| | oMM570 | GTTGAAGGAGATTATCAGCG |
| **pM1619** (*pMPS1-mps1(D580A),* i.e. *mps1*$^{D580A}$) *HIS3* SIV | oMM712 | GAAAGGTATCTTAAAAATCATTGCATTTGGTATAGCAAACGCGG |
| | oMM539 | GGCTTCTCAGAGCCATATCTAACG |
| **pM1620** (*pMPS1-mps1(Δ151-200),* i.e. *mps1*$^{\Delta 151-200}$) *HIS3* SIV | oMM713 | gcgacaaaatatgaaagaagatattacggcaaagtatgctgaaGAGGATTCTCACCAAACAAAC |
| | oMM570 | GTTGAAGGAGATTATCAGCG |
| **pM1621** (*pMPS1-mps1(Δ201-300),* i.e. *mps1*$^{\Delta 201-300}$) *HIS3* SIV | oMM714 | ccagccaataaatgcaagggagacagtggaattaccacttCCCAGGCGAAAAGTTTCTAC |
| | oMM570 | GTTGAAGGAGATTATCAGCG |
| **pM1642** (*pMPS1-mps1(L165E R170E),* i.e. *mps1*$^{L165E\ R170E}$) *HIS3* SIV | oMM803 | TGTCAAAGTCATCGCTTCCTTTGCAGGACCTTCCTTCGTTGTCCTATTGG |
| | oMM662 | CAGTTGTTTATAGAAGTAGC |

| | | |
|---|---|---|
| **pM1643** (*pMPS1-mps1(R151E R152E K154E R155E),* i.e. *mps1*$^{\text{R151E R152E K154E R155E}}$) *HIS3* SIV | oMM801 | TACGGCAAAGTATGCTGAAGAAGAAAGTGAGGAATTTTTAATATCCAATAGGACAACG |
| | oMM570 | GTTGAAGGAGATTATCAGCG |
| **pM1644** (*pMPS1-mps1(K169E R170E),* i.e. *mps1*$^{\text{K169E R170E}}$) *HIS3* SIV | oMM802 | TGTCAAAGTCATCGCTTCTTCTGCAGGACCCAGCTTCGTTG |
| | oMM662 | CAGTTGTTTATAGAAGTAGC |
| **pM1645** (*pMPS1-mps1(Δ4-50),* i.e. *mps*$^{\Delta\text{4-50}}$) *HIS3* SIV | oMM721 | caggtggtcgacggtatcgataagcttgatatcgatgtcaacaGAGATATTATCAAGTCATAATAATG |
| | oMM570 | GTTGAAGGAGATTATCAGCG |
| **pM1646** (*pMPS1-mps1(Δ151-157, 165-171),* i.e. *mps*$^{\Delta\text{151-157, 165-171}}$) *HIS3* SIV | oMM804 | gcgacaaaatatgaaagaagatattacggcaaagtatgctgaaATATCCAATAGGACAACGAAG |
| | oMM570 | GTTGAAGGAGATTATCAGCG |
| **pM1648** (*pMPS1-mps1(R151E R152E K154E R155E K169E R170E),* i.e. *mps1*$^{\text{R151E R152E K154E R155E K169E R170E}}$) *HIS3* SIV | oMM800 | GAAAGGAGAAGTAAGGAATTTTTAATATCCAATAGGACAACG |
| | oMM570 | GTTGAAGGAGATTATCAGCG |
| **pM1700** (*pGAL10-mps1(L165E R170E),* i.e. *mps1*$^{\text{L165E R170E}}$) | | Constructed via sub-cloning |
| **pM1701** (*pGAL10-mps1(R151E R152E K154E R155E),* i.e. *mps1*$^{\text{R151E R152E K154E R155E}}$) | | Constructed via sub-cloning |
| **pM1702** (*pGAL10-mps1(K169E R170E),* i.e. *mps1*$^{\text{K169E R170E}}$) | | Constructed via sub-cloning |
| **pM1703** (*pGAL10-mps1(Δ151-157, Δ165-171),* i.e. *mps1*$^{\Delta\text{151-157, }\Delta\text{165-171}}$) | | Constructed via sub-cloning |
| **pM1705** (*pGAL10-mps1 (R151E R152E K154E R155E R169E R170E),* i.e. *mps1*$^{\text{R151E R152E K154E R155E K169E R170E}}$) | | Constructed via sub-cloning |
| **pM1718** (*pMPS1-MPS1*, i.e. *MPS1*$^{\text{WT}}$) *URA3* SIV | | Constructed via sub-cloning |
| **pM1721** (*pGAL10-mps1(R155E L165E R170E),* i.e. *mps1*$^{\text{RLR>EEE}}$) | | Constructed via sub-cloning |
| **pM1722** (*pMPS1-mps1(R155E L165E R170E),* i.e. *mps1*$^{\text{RLR>EEE}}$) *HIS3 SIV* | | Constructed via sub-cloning |
| **pM1723** (*pMPS1-mps1(R155E L165E R170E),* i.e. *mps1*$^{\text{RLR>EEE}}$) *URA3* SIV | | Constructed via sub-cloning |
